# The biased adenosine-rich content of the HIV-1 genome serves as a molecular signature that facilitates efficient packaging

**DOI:** 10.1101/2025.05.01.651415

**Authors:** Hung R. Vuong, Qianzi Zhou, Sydney Lesko, Kasyap Tenneti, Keanu Davis, Shanyqua Scott, Moming Guo, Daphne Boodwa-Ko, Jenna E. Eschbach, Kamya Gopal, Jessica M. Porter, Qibo Wang, Ming Xia, Anthony Boateng, Yiqing Wang, Shawn Mohammed, Nakyung Lee, Alice Telesnitsky, Nathan Sherer, Sebla B. Kutluay

## Abstract

The HIV-1 genome (gRNA) has an unusually biased nucleotide content and is rich in adenosines. Selective packaging of the gRNA is thought to be driven by specific binding of the nucleocapsid (NC) domain of the viral Gag protein to the packaging signal (Ψ) in the host cell cytosol. However, deletion of regions within Ψ reduces—but does not completely abolish—genome packaging. To probe whether another feature of the gRNA may contribute to the selective gRNA packaging process, we replaced NC with heterologous RNA-binding domains (RBDs) with distinct RNA-binding properties. Surprisingly, despite disparate RNA binding specificities, all Gag-RBD chimeras successfully recruited the gRNA to the plasma membrane, suggesting that the initial gRNA recognition in the cytosol is not rate limiting. Notwithstanding, many chimeras exhibiting G/C binding specificity were arrested at the assembly stage. Only the Gag-SRSF5 chimera, which multimerized efficiently on adenosine-rich sequences on the gRNA, assembled efficiently and packaged gRNA at near wild-type levels. Importantly, rationally designed mutations that altered the A/G-rich binding specificity of Gag-SRSF5 decreased genome encapsidation efficiency. Furthermore, many Gag chimeras displayed potent dominant negative activities, highlighting NC functions as a targetable step in virus replication. Together, our findings reveal an unexpected aspect of the HIV-1 gRNA, its biased nucleotide content, as a key driver of selective genome packaging.

**SIGNIFICANCE:** How HIV-1 selectively packages its genome (gRNA) into virions is poorly understood. To probe this, we replaced the viral nucleocapsid (NC) protein with heterologous RNA-binding domains (RBDs) from cellular hnRNP and SR protein families. Remarkably, despite their distinct RNA-binding specificities, all Gag-RBD chimeras successfully recruited the gRNA to the plasma membrane, suggesting that the initial gRNA recognition is not rate limiting. Only the Gag-SRSF5 chimera packaged gRNA efficiently which correlated with its capacity to multimerize on adenosine-rich sequences on the gRNA. Notably, several Gag chimeras exhibited strong dominant negative effects, underscoring NC functions as a targetable step in virus replication. Together, these findings uncover that the biased nucleotide content of the HIV-1 gRNA facilitates its selective packaging into virus particles.

## INTRODUCTION

Selective packaging of viral genomes is essential for replication and contributes to rapid viral evolution. For HIV-1, the nucleocapsid (NC) domain of the major HIV-1 structural protein Gag mediates the selective encapsidation of a non-covalent dimer of the viral genomic RNA (gRNA) into assembling virions^1–5^. This process is highly efficient, with nearly all HIV-1 particles containing a dimer of gRNA^1^. How such remarkable enrichment is achieved despite competition from abundant cellular mRNAs in infected cells remains a long-standing question.

Efficient gRNA packaging is widely thought to be initiated in the cytosol through high affinity binding of NC to specific sites within the highly structured cis-acting packaging signal, Ψ, located near the 5’ end of the gRNA^6–11^. However, deletions within Ψ do not result in a complete loss of packaging^12–14^. Moreover, Gag binds with high affinity to both Ψ-containing and Ψ-null RNAs at physiological ionic strength^15^ and packages cellular mRNAs into virus particles in the absence of HIV-1 gRNA^16,17^. Hence, selective binding of Gag to Ψ is insufficient to account for selective gRNA packaging.

Although Gag-Ψ interactions initiate in the cytosol^10,18^, it remains unclear whether this early recognition event is essential for selective gRNA packaging. Live-cell imaging studies suggest that a small number of Gag molecules are needed to traffic and stably anchor the gRNA at the plasma membrane (PM)^19–21^, where they nucleate the recruitment of additional Gag molecules to form high-order Gag multimers and drive virus budding^3,22–24^. RNA also contributes structurally to virions^25^ and capsid (CA) mutations that disrupt Gag lattice assembly impair Ψ binding and genome packaging^26,27^. Together, these findings suggest that multiple Gag-binding sites within the gRNA increase local Gag concentration and promote Gag–Gag interactions during assembly^3,7^. However, the specific gRNA features that serve as a scaffold for Gag multimerization remain unknown.

The HIV-1 gRNA exhibits a strongly biased nucleotide content: rich in adenosines (∼36%) and poor in cytosines (∼18%) with purines overrepresented in all codon positions of all HIV-1 genes^28,29^. Herein, we tested whether this bias promotes Gag-gRNA interactions and facilitates selective gRNA packaging. To this end, we generated Gag chimeras with altered RNA binding specificities by replacing the NC domain with RNA binding domains (RBDs) from the heterogeneous nuclear ribonucleoprotein (hnRNP) and serine/arginine-rich splicing factor (SRSF) proteins. We hypothesized that chimeras with a preference for purine-rich sequences would efficiently package gRNA. Remarkably, despite disparate RNA binding specificities, all chimeras recruited the gRNA to the PM, suggesting that the initial gRNA recognition event is not rate limiting. However, only Gag-SRSF5, which preferentially binds A/G-rich sequences, packaged gRNA at near wild-type (WT) levels. Gag-SRSF5 efficiently multimerized on the HIV-1 gRNA at the PM and exhibited a purine-rich RNA binding specificity similar to WT Gag in immature virus particles. Mutations that reduced its A/G-rich binding specificity also impaired gRNA packaging. While all chimeras were non-infectious and many exhibited particle release defects, they co-assembled with WT Gag and potently inhibited virus infectivity. Together, these results indicate that cytosolic gRNA binding and PM trafficking alone are insufficient for selective gRNA packaging; rather, efficient Gag multimerization on A-rich HIV-1 gRNA at the PM drives preferential genome encapsidation over cellular RNAs.

## RESULTS

### Gag-SRSF5 chimera efficiently packages the HIV-1 gRNA into virus particles

We generated HIV-1_NL4-3_-based proviral clones in which the NC domain of Gag was replaced with tandem RBDs, including RNA-recognition motifs (RRMs) and K homology domains (KH), derived from members of the hnRNP and SRSF protein families. We reasoned that RBDs with a preference for binding to A-rich sequences might substitute for the genome packaging function of NC. The selected hnRNP and SRSF proteins were chosen for their established ability to bind to HIV-1 RNAs, but more importantly for their well-defined RNA binding specificities—hnRNP A1 recognizes “AGG” motifs; hnRNP F binds to G-rich oligonucleotides; hnRNP K binds to short C-rich tracks; SRSF1, 4 and 5 recognize “AAG” motifs ^30–35^. Controls included included replacement of NC with leucine zipper domain from GCN4 (bZIP) that facilitates Gag multimerization in the absence of NC ^36^ and complementation by NC itself (ΔNC-NC). All chimeras expressed similar levels of Gag and Gag-Pol as well as their processing intermediates in cells **(Fig. 1A).** Expression of the viral Env and Nef proteins, generated from spliced HIV-1 transcripts, was also comparable between WT Gag and Gag-RBD chimeric viruses, suggesting that the isolated RBDs did not alter viral mRNA splicing (**Fig. S1A**). Particle release was largely impaired for the Gag-hnRNP A1, -hnRNP F and hnRNP K chimeras **(Fig. 1A)**, whereas Gag-SRSF1 and Gag-SRSF4 exhibited more modest release defects, and Gag-SRSF5 release was largely unaffected **(Fig. 1A)**. Gag-SRSF5 particles contained levels of Gag-Pol and integrase comparable to WT Gag and ΔNC-NC virions, indicating normal Gag-Pol incorporation and processing **(Fig. 1A)**. However, given the multiple functions of NC in HIV-1 replication beyond packaging^37–40^, all chimeric viruses including Gag-SRSF5 were non-infectious (**Fig. S1B**).

**Figure 1.**
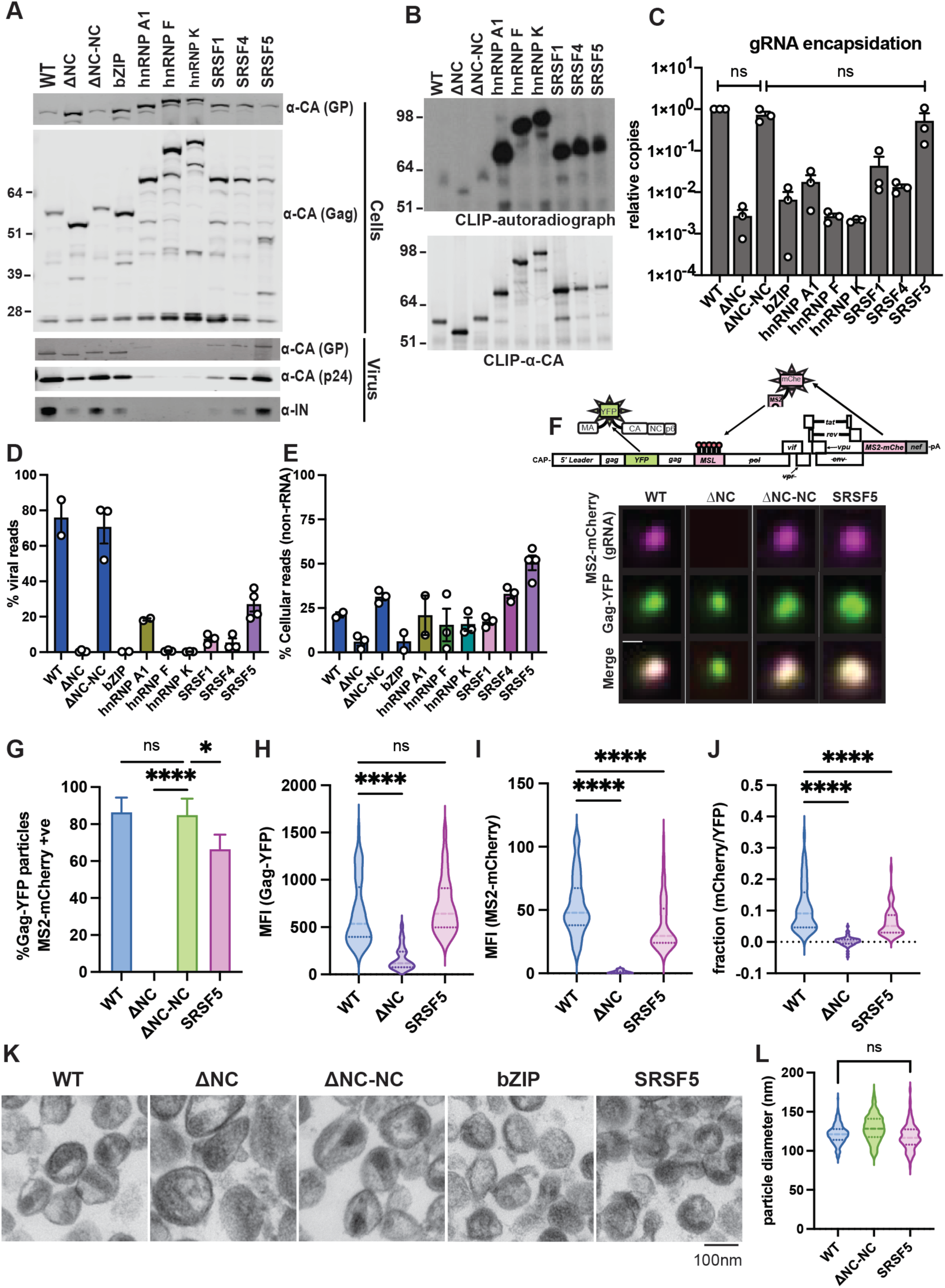
Gag-SRSF5 chimera efficiently packages HIV-1 gRNA. **A.** HEK293T cells were transfected with HIV-1_NL4-3_ proviral plasmids encoding the indicated Gag-RBD chimeras. Cell lysates and purified virions were analyzed with immunoblotting for Gag-Pol (GP), Gag, Capsid (p24) and Integrase (IN). **B.** Representative PAR-CLIP autoradiogram of Gag:RNA adducts (top) and corresponding anti-HA immunoblot of Gag proteins (bottom) immunoprecipitated from cells that were transfected with protease null proviral plasmids encoding the indicated Gag-RBD chimeras. **C.** RT-qPCR quantification of relative copies of gRNA from purified Gag-RBD chimeric virus particles (WT set to 1). Data show the average of 3 independent experiments, error bars represent standard error of the mean (SEM). Statistical analyses were done by one-way ANOVA with Dunnet’s multiple comparison test for correction, whereby n.s. denotes non-significant comparisons. Other group comparisons were *P*<0.01 but are not displayed due to clarity. **D-E.** Proportion of reads that map to viral RNA (D) and cellular non-ribosomal RNAs (E) obtained from total RNA-seq experiments performed on purified virions. Data show the average of 2-3 independent experiments, error bars represent the SEM. **F.** Representative fluorescence microscopy images of HIV-1 particles harvested from 293T cells generating two-color viruses encoding a Gag chimera-YFP fusion protein and gRNAs engineered to carry MS2 RNA stem loops (MSL) that are tightly bound by self-encoded MS2-mCherry tracker proteins. Cartoon depiction of the two-color (Gag-YFP/MS2-mCherry) reporter virus is depicted above the images. **G.** Percentage of Gag-YFP+ particles that contain gRNA (mCherry^+^YFP^+^) based on single-virion analysis is shown. Data show the mean from 3 independent experiments, error bars show the standard deviation. **H-J**. Violin plots showing Gag-YFP (H), MS2-mCherry (I) and MS2-mCherry/Gag-YFP (J) mean fluorescence intensity (MFI) ratios measured for WT, Gag-ΔNC, and Gag-SRSF5 particles shown in panel F. 60 particles were quantified for each condition. **K.** Representative thin section electron microscopy (TEM) images showing WT Gag, Gag-ΔNC, Gag-ΔNC-NC, Gag-bZIP, and Gag-SRSF5 virions. Scale bar = 100 nm. **L.** Violin plot showing the particle diameters (n=200) obtained from panel K. For panels G-L \**P*<0.05; **** *P*<0.0001; n.s. non-significant by one-way ANOVA with Dunnet’s multiple comparison test for correction.

As intended, all Gag-RBD chimeras bound to RNAs in cells and formed Gag-RNA adducts, with each chimera associating with total RNA at levels significantly higher than WT Gag (**Fig. 1B, S1C**). However, RT-qPCR analysis of purified virus particles revealed that only Gag-SRSF5 packaged gRNA with an efficiency resembling WT Gag **(Fig. 1C)**. Interestingly, host 7SL RNA which is known to be enriched in HIV-1 virions^10,41–43^, was also packaged at comparable levels to WT Gag in Gag-SRSF5 particles (**Fig. S1D**). On the other hand, Gag-SRSF5 appeared to be less selective for gRNA compared to WT Gag and encapsidated a higher proportion of multiply spliced HIV-1 RNAs (**Fig. S1D, S1E**), possibly in part due to its known splicing regulatory functions. In agreement, RNA-seq analysis of virion-incorporated RNAs further showed that ∼30% of RNAs in Gag-SRSF5 particles were HIV-1-derived **(Fig. 1D)**, while cellular non-rRNAs—predominantly composed of mRNAs and 7SL RNA—were enriched approximately twofold compared to WT Gag and ΔNC-NC virions **(Fig. 1E, S1F)**. Northern blotting both confirmed these results and demonstrated that gRNA and 7SL RNA packaged in Gag-SRSF5 particles were intact (**Fig. S1G**). Among the remaining chimeras, Gag-hnRNP A1 was the next most efficient at gRNA packaging after taking into account the reduced number of released virus particles (**Fig. S1E**), with approximately 20% of RNAs in Gag-hnRNP A1 virus particles HIV-1-derived (**Fig. 1D**).

Single-virion imaging using YFP-tagged Gag and HIV-1 gRNA labeled using the bacteriophage-derived MS2 system^19,44–46^ showed that ∼70% of Gag-SRSF5 particles encapsidated gRNA **(Fig. 1F, 1G).** Though we didn’t directly assess whether the packaged genomes are monomers or dimers, gRNA signal intensity per virion trended lower for Gag-SRSF5 particles relative to WT Gag and Gag-ΔNC-NC (**Fig. 1H-J**), a result consistent with the hypothesis that not all Gag-SRSF5 particles were packaging gRNA as dimers. Expectedly^1^, ∼90% of WT and ΔNC-NC virions contained gRNA, whereas most ΔNC particles lacked detectable genomes (**Fig. 1F, 1G**). Virion-associated Gag-YFP levels were comparable between WT and Gag–SRSF5 particles (**Fig. 1H**), whereas ΔNC particles incorporated substantially less Gag-YFP (**Fig. 1H**), consistent with impaired assembly.

To further examine the morphology of Gag-SRSF5 particles, we performed thin-section transmission electron microscopy (TEM) of purified virions. Like WT and ΔNC-NC particles, Gag-SRSF5 particles were ∼100nm in diameter but contained eccentrically localized condensates or displayed immature morphology **(Fig. 1K, 1L).** Biochemical fractionation of viral cores from Gag-SRSF5 particles revealed unstable or improperly assembled cores, similar to those observed for ΔNC and bZIP viruses, likely due to the presence of aberrant Gag processing intermediates (**Fig. S1H-J**). These defects correlated with early reverse transcription defects of Gag-SRSF5 particles in target cells (**Fig. S1K**) and a corresponding reduction in virion infectivity (**Fig. S1B**).

### Gag-RBD chimeras efficiently recruit HIV-1 gRNA to the PM but many display assembly defects

The release defects of the several Gag-RBD chimeras prompted us to evaluate at what step of virus assembly they were blocked at, which we reasoned would help explain what makes WT Gag and Gag-SRSF5 successful at gRNA packaging. Like WT Gag, all Gag-RBD chimeras localized in the cytosol and at the PM at steady state **(Fig. 2A, S2A, S2B)**. The control Gag myristoylation mutant (G2A) which is defective in PM anchoring and was enriched in the cytosol fraction (**Fig. 2A, S2A**). Surprisingly, all Gag-RBD chimeras recruited gRNA to the PM as efficiently as WT Gag, with hnRNP A1 exhibiting markedly higher levels **(Fig. 2B)**. Notably, Gag-hnRNP A1 preferentially trafficked gRNA to the PM relative to multiply spliced viral RNAs, whereas other chimeras, including Gag-SRSF5, were comparably less selective **(Fig. 2B)**. In contrast, the abundance of the control host GAPDH mRNA at the PM was independent of the presence of Gag with a functional NC domain or RBDs (**Fig. 2B**). Single cell imaging of dually labeled viruses confirmed that all Gag chimeras efficiently recruited the gRNA to the PM **(Fig. 2C, 2D, S2C, S2D)**. However, only Gag-SRSF5 formed WT-like foci with gRNA at the PM whereas the other Gag chimeras often formed sheet-like assemblies at the PM consistent with their assembly defects **(Fig. 2D, S2C, S2D)**. We also noted that a fraction of Gag-hnRNP F displayed strong nuclear localization in many cells (**Fig. 2D**), potentially mediated by the central glycine-tyrosine-arginine-rich (GYR) domain that lies between hnRNP F RRM2 and RRM3 and functions as a nuclear localization signal^47^.

**Figure 2.**
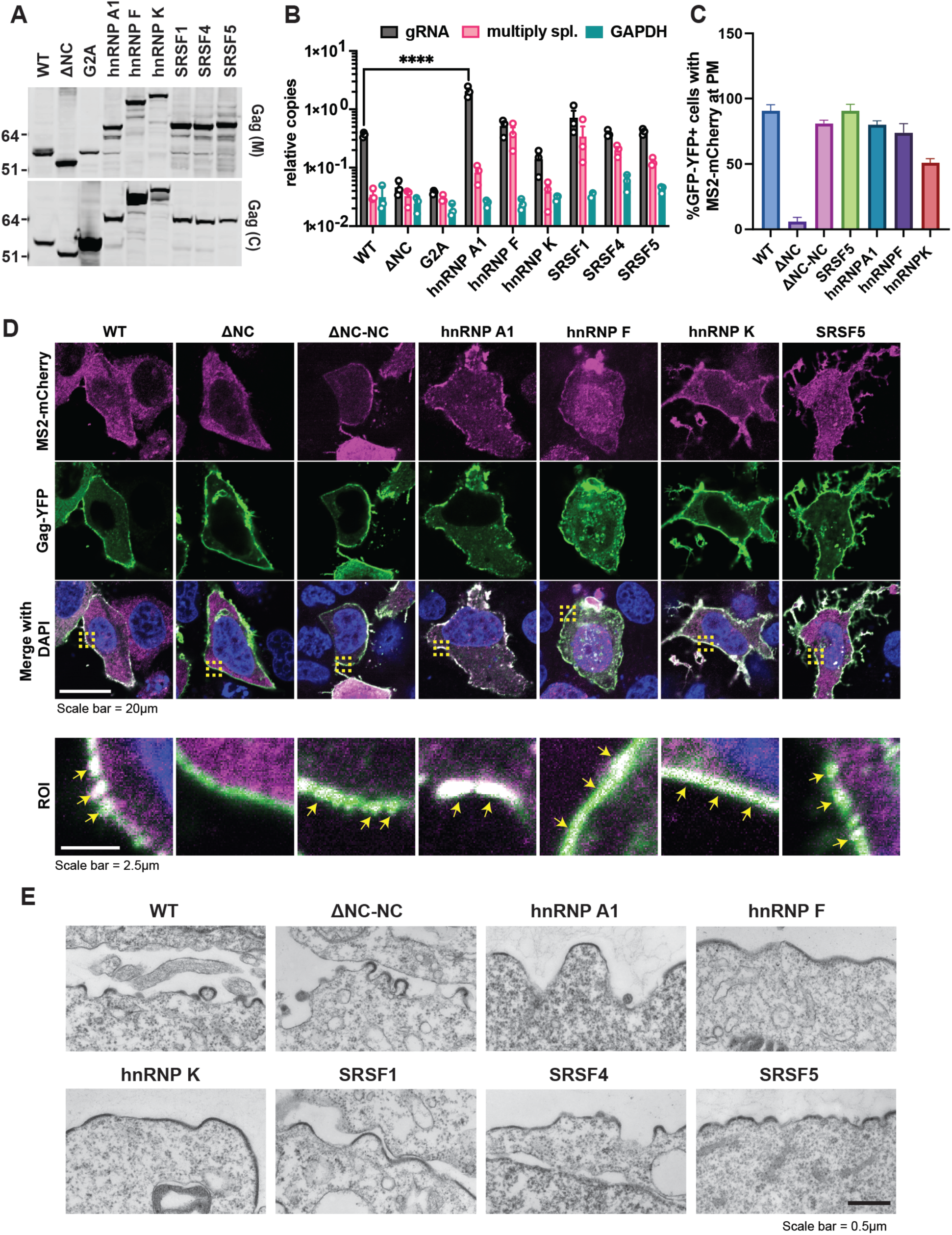
Gag-RBD chimeras efficiently recruit and retain gRNA at the PM. **A.** Immunoblot analysis of Gag from membrane (M) and cytoplasmic (C) fractions isolated from 293T cells transfected with the indicated proviral (PR null) Gag-RBD plasmids using a mouse monoclonal anti-HA antibody. **B.** RT-qPCR quantitation of the relative PM/cytosol ratios of gRNA, multiply spliced HIV-1 mRNA, and GAPDH mRNA copies. Data show the average of three independent experiments, error bars represent SEM. **** *P*<0.0001 by one-way ANOVA with Dunnet’s correction. Non-significant comparisons are not displayed for clarity. **C.** Quantification of Gag-gRNA colocalization at the PM from fluorescence imaging of HeLa cells generating the two-color (Gag-YFP/MS2-mCherry) reporter viruses described in Fig. 1F modified to encode the indicated Gag-RBD chimera. **D.** Representative fluorescence microscopy images of HeLa cells generating the two-color (Gag-YFP/MS2-mCherry) reporter viruses. . Scale bars represent 20 μm in full images and 2.5 μm in regions of interest (ROI) at the PM. ROIs are indicated by yellow dotted boxes. Yellow arrowheads indicate where Gag and gRNA colocalize. **E.** Representative TEM images from two independent experiments showing virion assembly sites at the PM of 293T cells transfected with the indicated Gag plasmids. Scale bar represents 500 nm.

To interrogate whether Gag-RBD chimeras defective for assembly could multimerize at the PM but not generate the membrane curvature necessary for ESCRT-III recruitment^48^, we conducted TEM analyses in virus producer cells. Consistent with results from our fluorescence imaging studies (**Fig. 2D, S2D**), Gag-RBD chimeras with release defects formed extended electron-dense sheet-like structures at the PM **(Fig. 2E)**, indicating that they were capable of multimerization and lattice formation. However, unlike WT Gag and ΔNC-NC, which formed well-defined curved buds, the electron dense sheet-like structures from hnRNP A1, F, K and Gag-SRSF1 chimeras had little to no membrane curvature. In contrast, Gag–SRSF4 and Gag–SRSF5 generated buds more similar in size and morphology to WT and ΔNC-NC controls **(Fig. 2E)**. Altogether, these data demonstrate that Gag-RBD chimeras possessing diverse RNA-binding modules can recruit the gRNA to the PM. However, defects in post-recruitment events impair membrane curvature, budding, and efficient gRNA packaging for most chimeras.

### Gag-RBD chimeras exhibiting diverse RNA-binding specificities bind similarly to HIV-1 gRNA in cells

To identify the feature of Gag-SRSF5 that allows it to package gRNA almost as efficiently as WT Gag, we examined the RNA-binding specificities of Gag-RBD chimeras using PAR-CLIP at different stages of virus particle assembly. In cells, similar to WT Gag, only a small fraction of RNAs bound by most Gag–RBD chimeras were HIV-1–derived (**Fig. S3A**). This indicates that the ability of Gag-SRSF5 to package gRNA is not due its association with higher levels of gRNA in the cytosol. In contrast, Gag-hnRNP A1 consistently bound 5-10-fold more HIV-1 RNAs than WT Gag (**Fig. S3A**), consistent with its enhanced ability to recruit gRNA to the PM (**Fig. 2B**). Also similar to WT Gag, most Gag chimeras—except Gag-hnRNP F— were heavily associated with cellular tRNAs (via the matrix domain¹¹) and mRNAs (**Fig. 3A**); with mRNAs constituting a larger fraction of cell-derived RNAs bound by the Gag-hnRNP A1, -SRSF4, and -SRSF5 chimeras (**Fig. 3A**). WT Gag and all chimeras also bound to introns, possibly reflecting a subpopulation of Gag accumulating in the nucleus^49–51^. Nuclear trafficking of Gag-hnRNP F (**Fig. 2D**) could explain why tRNAs represented a smaller fraction of reads relative to intron-mapping reads for this chimera (**Fig. 3A**), as well as hnRNP F’s known strong association with introns^33,35,47^. Motif analysis of bound mRNAs showed that the sequence preferences of Gag–RBD chimeras aligned with the known RNA-binding specificities of their respective hnRNP/SR protein–derived RBDs **(Fig. S3B)**^33,35,52–55^.

**Figure 3.**
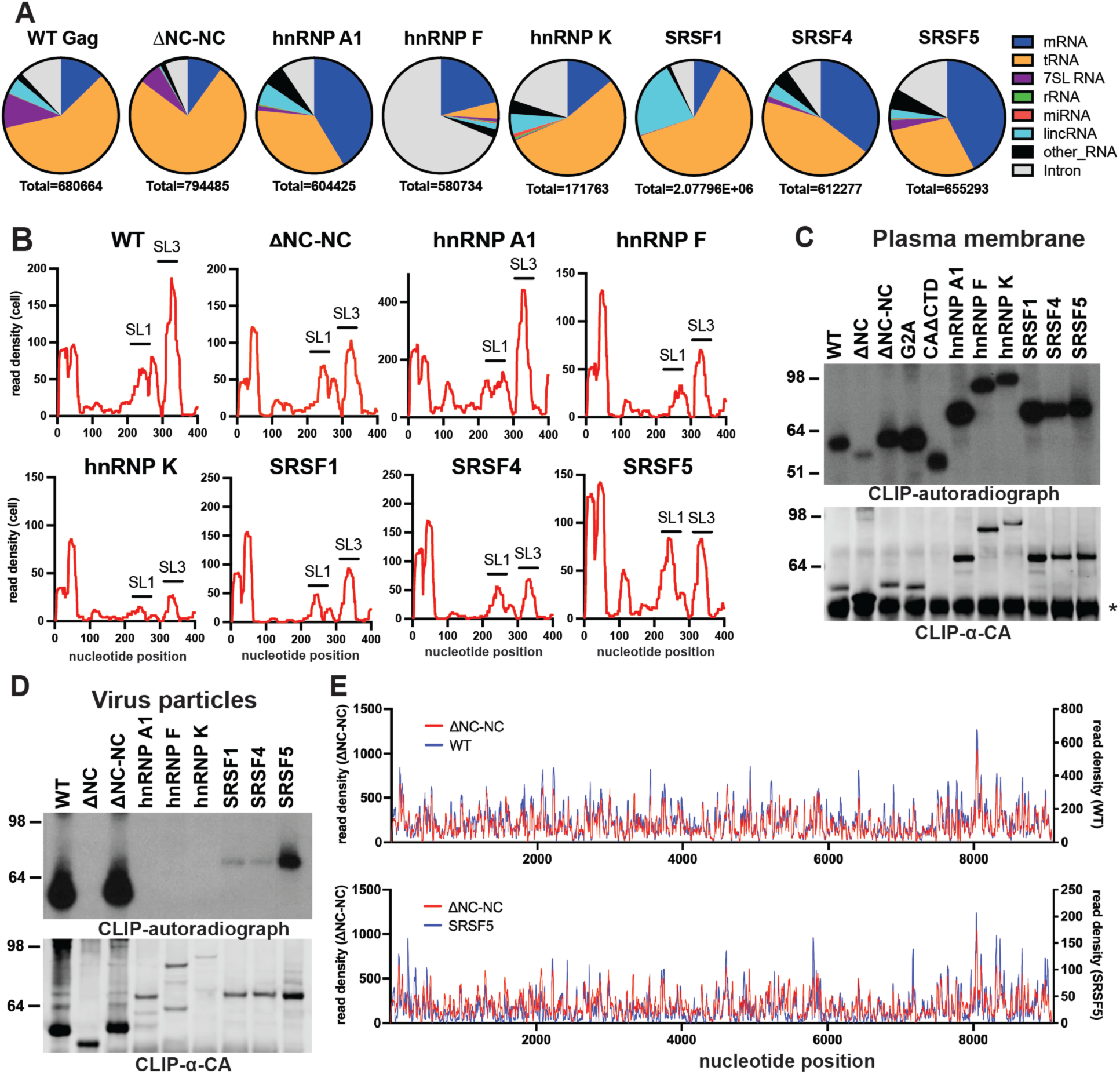
CLIP-seq analysis of Gag-RBD chimeras reveals the unique ability of Gag-SRSF5 to multimerize on the gRNA in virus particles. **A.** Classification of read clusters that map to cellular RNAs generated from Gag-CLIP experiments done in cell lysates; total number of reads is indicated below each pie chart. **B.** Representative read density distribution within the 5’ leader (1-400 nt, x-axis) from Gag-RBD CLIP experiments done in cell lysates. Location of stem loops 1 and 3 (SL1, SL3) are indicated by black lines. **C-D.** Representative autoradiograms of Gag-RNA complexes (top) and immunoblot analysis of Gag proteins (bottom) immunoprecipitated from the PM fraction (C) or immature virus particles (D) in CLIP experiments. (*) marks the IgG-heavy chain from the immunoprecipitating anti-HA antibody. **E.** Read density distribution on full-length HIV-1 gRNA (x-axis) from WT Gag, Gag-ΔNC-NC and Gag-SRSF5 CLIP-seq experiments done in immature virus particles.

On the HIV-1 genome, WT Gag and Gag-ΔNC-NC bound to discrete sites within the 5’ leader, including SL1 and SL3 (**Fig. 3B, S3C**), consistent with prior reports^10^. Surprisingly, all Gag-RBD chimeras also preferentially bound these sites despite their diverse RNA-binding specificities, though the extent with which they were bound to SL1 and SL3 varied (**Fig. 3B, S3C**). We also tested whether progressive deletions of structured RNA elements within the 5’ leader would affect the efficiency of gRNA packaging by Gag-SRSF5 (**Fig. S3D**), observing only modest (2-fold) relative reductions in packaging for a subset of mutations (**Fig. S3E, S3F**). We interpreted these data as consistent with the hypothesis that features outside of the 5’ leader are also contributing significantly to the efficiency of genome packaging.

### Gag-SRSF5 multimerizes on the A-rich HIV-1 gRNA sequences

Prior CLIP studies have established a correlation between Gag multimerization during virion assembly and activation of a switch in Gag RNA-binding specificity; from G-rich sequences towards A-rich motifs ^10^. To determine if a similar specificity switch underpinned Gag-SRSF5’s ability to packaging gRNA similar to WT Gag, we performed comparative CLIP experiments for all Gag chimeras from isolated PM fractions and immature virus particles. In membrane fractions, WT Gag and Gag-RBD chimeras all bound higher overall levels of RNA relative to in the cytoplasm (**Fig. 1B, S1C** vs. **Fig. 3C, S4A)**, consistent with enhanced engagement with cognate RNAs upon multimerization at the PM during the onset of virus particle assembly. In contrast, in virions, only Gag-SRSF5 yielded levels of protein:RNA complexes similar to WT Gag, whereas Gag-SRSF1 and Gag-SRSF4 were largely impaired in RNA-binding **(Fig. 3D, S4B)**. Expectedly, viral RNA-derived CLIP reads were enriched in virions relative to cells for WT Gag **(Fig. S4C**), but relatively low overall for Gag-SRSF5 **(Fig. S4C)**, consistent with its less selective gRNA packaging (e.g. **Fig. 1G**).

Strikingly, in immature virus particles, the Gag-SRSF5 binding profile on gRNA closely mirrored that of WT Gag and Gag-ΔNC-NC (**Fig. 3E**), with a statistically significant correlation nearing that between WT Gag vs. Gag-ΔNC-NC (**Fig. S4D**). In virus particles, Gag-SRSF5-bound host mRNAs also resembled those bound by WT Gag and Gag-ΔNC-NC, with mRNAs, intron-containing RNA, and 7SL RNA representing the most frequently bound RNAs (**Fig. S4E).** Moreover, A/G-rich motifs were enriched in cellular mRNAs bound by Gag-SRSF5, WT Gag, and Gag-ΔNC-NC in virus particles **(Fig. S4F)**. In contrast to the genome binding profiles for WT Gag and all Gag chimeras in the cytoplasm **(Fig. S3C)**, WT Gag and Gag-SRSF5 was associated with a larger number of sites throughout the HIV-1 gRNA at the PM and less prominently at the 5’ UTR **(Fig. S4G),** consistent with a switch in binding specificity to A-rich sequences during the onset of virus particle assembly. Consistent with this model, mutation of Gag to reduce multimerization at the PM (Gag-CAΔCTD) markedly restricted Gag’s capacity to bind across A-rich regions of the gRNA (**Fig. S4G**). Gag-hnRNP A1 and Gag-hnRNP F, which bind to G-rich sequences in cells, similarly showed little to no increase in A-rich binding sites at the PM (**Fig. S3C vs. Fig. S4G**). Taken together, these results provided strong support for the notion that the abilities of Gag and Gag-SRSF5 to efficiently multimerize on the A-rich gRNA may underlie their similar capacities to efficiently package the HIV-1 genome.

### RNA binding properties of Gag-SRSF5 variants with reduced gRNA packaging efficiency

To more directly test whether Gag-SRSF5’s preference towards adenosine-rich motifs is needed to package gRNA, we introduced mutations within the SRSF5 RRMs rationally designed to reduce purine binding. To this end, we targeted two tandem RRMs bridged by a flexible linker in SRSF5, predicted to mediate recognition of a minimal GGA motif based on a prior structural study of SRSF1’s RRM2 domain^56^. In this model, SRSF1 RRM2 residues D136, K138 and D139 were shown to serve as the basis for the extensive hydrogen binding network that defines the purine-rich (GGA)-specific recognition event. Guided by this model, we designed two Gag-SRSF5 mutants, mGGA-1 and mGGA-2, carrying substitutions in the homologous residues (e.g. D123, K125, D126), predicted to weaken GGA-specific RNA recognition (**Fig. S5A**). In addition, we engineered mutants predicted to shift the RNA-binding specificity of SRSF5 towards a CCG motif (mCCG-1 and mCCG-2) or a GGG motif (mGGG) (**Fig. S5A**).

Reassuringly, these mutations did not impair the overall ability of Gag-SRSF5 to bind RNA in cells, and all Gag-SRSF5 variants immunoprecipitated with RNA at higher than WT levels **(Fig. S5B, S5C)**. In contrast, in virus particles, the variants bound to less RNA than parental Gag-SRSF5 **(Fig. S5D, S5E)**. Consistent with our expectations, CLIP analysis showed that binding sites for Gag-SRSF5 mCCG-2 and mGGG mutants in cellular RNAs were less adenosine-rich and more cytosine-rich (**Fig. S5F, S5G**), while uracil and guanosine frequencies were unchanged (**Fig. S5H, S5I**). Because 4SU-based crosslinking may underestimate shifts in specificity, we repeated CLIP using 6-thioguanosine. Consistent with altered specificity, Gag-SRSF5 mCCG-2 and mGGG mutants crosslinked to higher levels of RNA (**Fig. S5J**). Altogether, the above data support the intended shift toward C-rich or G-rich sequence recognition **(Fig. S5G, S5J)** and away from A-rich sequence recognition **(Fig. S5F)**.

Gag-SRSF5 mGGA-1 and mGGA-2 mutants packaged approximately two-fold lower levels of gRNA, multiply spliced HIV-1 RNAs, and 7SL RNA relative to the parental Gag-SRSF5 control, though this difference was not deemed statistically significant for gRNA **(Fig. 4A-C)**. The Gag-SRSF5 mCCG-1, mCCG-2 and mGGG mutants also exhibited stronger defects in packaging of gRNA, multiply spliced viral RNAs, and 7SL RNA **(Fig. 4A-C)**. As additional specificity controls, packaging of cellular RPL36A, RPS6 and RPS16 mRNAs that we and others have found to be abundantly incorporated into HIV-1 VLPs^16^ was largely unaffected **(Fig. S5K-M)**.

**Figure 4.**
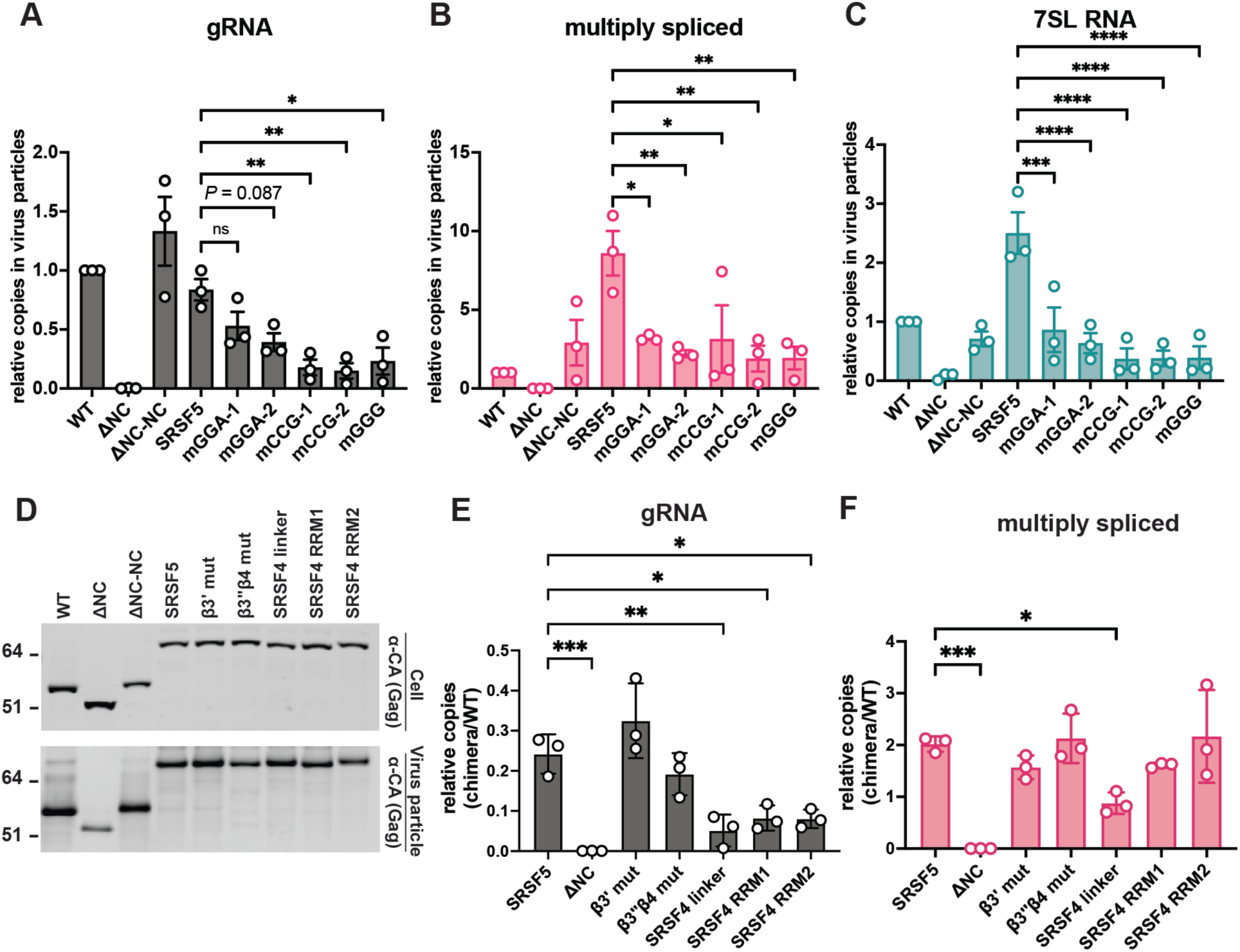
RNA binding properties of Gag-SRSF5 variants with impaired gRNA packaging. A-C. RT-qPCR quantification of relative copies of gRNA, multiply spliced HIV-1 RNA, and 7SL RNA from HIV-1 particles collected from 293T cells expressing the indicated Gag. Data show the average of 3 independent experiments and the error bars represent SEM. **D.** HEK293T cells were transfected with proviral plasmids (PR null) encoding the indicated Gag-SRSF5/SRSF4 chimeric proteins. Cell lysates and purified virus particles were harvested 2 days post-transfection and analyzed with immunoblotting for Gag. **E-F.** RT-qPCR quantification of relative copies of gRNA (E) and multiply spliced HIV-1 RNAs (F) from HIV-1 particles collected from 293T cells expressing the indicated Gag. Data show the average of 3 independent experiments and the error bars represent SEM. (\**P* < 0.05; \*\**P* < 0.01; \*\*\**P* < 0.001; \*\*\*\**P* < 0.0001 by one-way ANOVA with Dunnet’s multiple comparison test for correction, whereby n.s. denotes non-significant comparisons). Some n.s. comparisons are not displayed due to clarity.

We next determined the features of SRSF5 that enabled it to package gRNA efficiently compared to the closely related SRSF4 protein, despite these proteins’ similar reported sequence binding specificities. We noted that SRSF4 RRM2 β3’ and β3’’β4 sheets near the RNA binding pocket were more electronegative, and therefore introduced the respective SRSF4 substitutions into the Gag-SRSF5 backbone (β3’ mut and β3’’β4 mut). In addition, we replaced the linker, RRM1, and RRM2 of Gag-SRSF5 with that of SRSF4. All mutants were well expressed and supported efficient particle release **(Fig. 4D)**. While Gag-SRSF5 β3’ mut and β3’’β4 mut variants packaged gRNA at levels equivalent to Gag-SRSF5, chimeras containing the SRSF4 linker, RRM1, or RRM2 displayed significantly reduced gRNA packaging **(Fig. 4E)**. Only the SRSF4 linker mutant reduced packaging of multiply spliced viral RNAs **(Fig. 4F)**, indicating a more specific effect on gRNA packaging for the SRSF4 RRM/linker swaps. Together, these findings further supported the conclusion that A-rich RNA binding facilitates gRNA packaging and suggested that unique features within the SRSF5 RRM and linker domains regulate Gag-SRSF5’s selective packaging ability of the HIV-1 gRNA.

### Gag-RBD chimeras potently interfere with virion infectivity

Given the release defects of Gag-RBD chimeras and their potential relevance to antiviral strategies, we next tested their capacity to inhibit HIV-1 replication when expressed in *trans*. In co-transfected cells, WT Gag enhanced but didn’t fully restore virion incorporation of Gag-hnRNP A1, F and K with comparably small effects on Gag-SRSF1, 4 and 5 **(Fig. 5A, S6A)**. Co-assembly of Gag-hnRNP A1, Gag-hnRNP K, Gag-SRSF1, Gag-SRSF4, and Gag-SRSF5 with WT Gag reduced viral titers by ≥100 fold, with negligible effects on particle release **(Fig. 5B, S6B)**. Except for Gag-hnRNPK, which reduced gRNA packaging by ∼3-fold, Gag-RBD chimeras did not interfere with gRNA packaging and Gag-SRSF5 modestly increased the amount of packaged gRNA (**Fig. S6C**). Thus, the potent dominant negative effects of Gag-RBD chimeras were unlikely to result from interference with gRNA packaging *per se*. Notably, Gag-hnRNP K exerted the strongest dominant negative effect despite low levels of incorporation into particles in comparison to WT Gag **(Fig. 5A, 5B)**, with these effects confirmed as dose-dependent **(Fig. 5C, S6D)**. Interestingly, co-transfection with Gag-ΔNC and Gag-bZIP also led to a significant decrease in infectivity (**Fig. 5B**), suggesting that the number of NC copies per virion is finely tuned.

**Figure 5.**
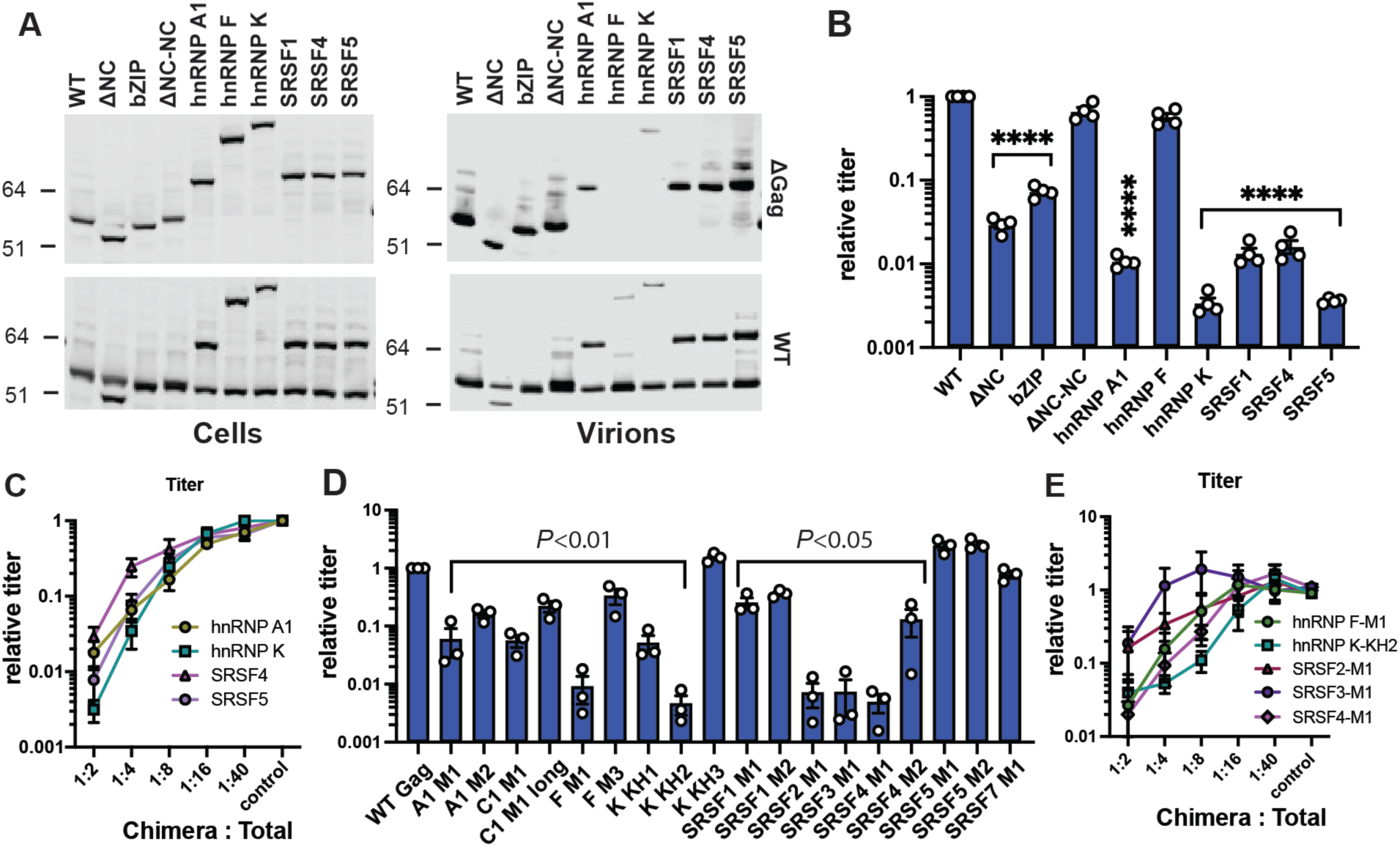
Gag-RBD chimeras are potently dominant negative. **A.** Proviral plasmids encoding ΔGag or WT Gag was co-transfected with the indicated Gag-RBD proviral plasmids in 293T cells. 2 days post-transfection, cell lysates and purified virions were harvested for immunoblot analysis of Gag using a mouse monoclonal antibody against the HA-tag. **B.** 293T cells were transfected with equal ratio of Gag-RBD chimeric HIV-1 proviral plasmids and a WT Gag HIV-1 proviral plasmid. Progeny virions were titered on TZM-bl reporter cells, with dextran-sulfate added to limit infection to one cycle. The titers were presented as relative to WT which was set to 1. **C.** Infectious titers of viruses released from 293T cells co-transfected with a WT Gag proviral plasmid and variable amounts of the Gag-RBD chimeras. **D.** 293T cells were transfected as in Panel B but with the indicated Gag-RBD chimeras bearing single RRMs. Mean infectious titers of released viruses on TZM-bl indicator cells is shown. **E.** Infectious titers of viruses released from 293T cells co-transfected with a WT Gag proviral plasmid and variable amounts of the single domain Gag-RBD chimeras. Data show the mean of four (panel B, C) and three (panel D, E) independent replicates with error bars displaying the SEM. ****P<0.0001 by one-way ANOVA with Dunnet’s correction. Non-significant comparisons are not displayed for clarity.

To further map the dominant negative activities, we next generated Gag-RBD chimeras encoding each single RRM from hnRNP A1, F, K, or SRSF 1, 4, and 5 **(Fig. S7A)**, and further screened RRMs derived from other hnRNP/SR proteins that naturally bear a single RBD (e.g. hnRNP C, SRSF2, SRSF3, SRSF7). All Gag single-RBD chimeras generated virus particles with release efficiencies that largely correlated with Gag-RBD chimera expression level in cells **(Fig. S7B),** but these particles were universally non-infectious **(Fig. S7C)**. Interestingly, while all Gag chimeras bound RNA in cells **(Fig. S7B),** none—including the Gag-SRSF5-RRM1 and Gag-SRSF5-RRM2 chimeras—could package gRNA **(Fig. S7CD**. In competition assays, most but not all chimeras reduced the infectivity of WT virions without impairing particle release (**Fig. 5D, S7E**), with unique patterns emerging. For example, the individual RRMs of hnRNP A1 reduced infectivity by 10-fold (**Fig. S7E**) as compared to the ∼100-fold reduction previously observed for these RRMs in tandem (**Fig. 5B, 5D**), suggesting cooperativity. For the Gag-hnRNP K and Gag-SRSF4 chimeras, the bulk of their inhibitory activity could be mapped to their KH2 and RRM1 domains, respectively (**Fig. S7E**). Interestingly, while the full RBD Gag-hnRNP F chimera had no effect on infectivity (**Fig. 5B, 5D**), the Gag-hnRNP F-RRM1 chimera was potently dominant negative (**Fig. S7E**). Of the proteins with single RBDs, Gag-SRSF2 and Gag-SRSF3 were the most potent, whereas Gag-SRSF7 did not inhibit infectivity and Gag-hnRNP C1 was only modestly inhibitory (**Fig. 5D**). With the exception of the Gag-SRSF3-RRM1 chimera, that decreased gRNA packaging by 5-fold, none of the chimeras interfered with WT Gag’s ability to package gRNA (**Fig. S7F**), again suggesting effects on virion structural integrity. Dose-response analysis further identified Gag-hnRNP K-KH2 as the most potent inhibitor **(Fig. 5E, S7G)**. Although the precise mechanisms by which Gag-RBD chimeras inhibit infectivity remain to be determined, these results demonstrate that perturbing the precision of NC-RNA recognition can potently inhibit HIV-1 replication even downstream of successful gRNA packaging.

## DISCUSSION

Our findings challenge the prevailing model that NC-mediated recognition of the 5′-UTR in the cytosol is the primary determinant of HIV-1 genome packaging. We show that heterologous RBDs with diverse RNA-binding specificities can substitute for NC to recruit gRNA to the PM, albeit with varying selectivity (**Fig. 2B**). Despite successful recruitment, only Gag-SRSF5 packaged gRNA at near WT levels which correlated with its unique ability to multimerize on the A-rich genome at the PM and in immature virus particles. Together, these results indicate that PM recruitment of gRNA alone is insufficient and that additional post-recruitment events facilitate efficient genome packaging.

An unexpected finding from our study is that Gag-RBD chimeras with diverse RNA-binding specificities bound regions within Ψ in cells, while still maintaining their established RNA-binding preferences for cellular RNA targets. Multiple studies have shown that WT Gag recognizes Ψ via high affinity interactions with unpaired guanosines within SL1 and SL3^11,57–59^. Because the Gag-RBD chimeras tested here, with the exception of Gag-hnRNP K, preferentially bind to single-stranded G-rich or A/G-rich sequences, it is possible that they engaged Ψ through similar interactions with unpaired guanosines. Moving forward, it will be interesting to determine whether RNA-binding modules that recognize dsRNAs can similarly recruit the HIV-1 gRNA to the PM and successfully replace NC. In addition, understanding whether Ψ and HIV-1 gRNA bear unique structural features that more readily destine the gRNA for PM trafficking by diverse RBDs will provide crucial insight into selective gRNA packaging.

The well-defined specificity of the hnRNP and SRSF proteins studied herein allowed us to obtain direct evidence to support the notion that A/G-rich binding by Gag-SRSF5 facilitates efficient HIV-1 genome packaging. It is, however, intriguing why the closely related Gag-SRSF1 and Gag-SRSF4 chimeras with similar RNA-binding properties failed to package gRNA. We propose that this selectivity does not necessarily argue against the putative role of A/G-rich binding in efficient genome packaging, because additional factors likely differ among these proteins. For example, Gag-SRSF1 and -SRSF4 are bound to less RNA in virions than Gag-SRSF5 (**Fig. 3D**), and in cells only one of their two RRMs (RRM1) bound RNA robustly, whereas both RRMs of SRSF5 were engaged in strong RNA binding (**Fig. S7A**). Consistent with weak binding, neither of the two SRSF4 RRMs could substitute for SRSF5 RRMs (**Fig. 4E**). Moreover, the individual SRSF5 RRMs were insufficient for gRNA packaging (**Fig. S7C**), also consistent with the notion that both RRMs of SRSF5 bear unique features and functioned cooperatively.

The finding that gRNA encapsidation efficiency decreased when the electropositive RRM linker of SRSF5 was replaced with the more electronegative linker of SRSF4 also suggests that electrostatic interactions contribute to gRNA packaging (**Fig. 4E**). This aligns with prior work wherein basic residues within the RNA-binding domains from the HIV-1 Rev protein or the herpes simplex type 1 ICP27 protein were found successfully substitute for NC residues in the Rous sarcoma virus Gag NC protein ^60^. The modular comparative strategy based on Gag-RBD chimeras developed for this study should provide additional opportunities for further dissection of the key structure-function relationships that underpin the differential control of NC-gRNA interactions in the cytoplasm and at the PM.

Beyond providing a unique tool for studying gRNA packaging, our Gag-RBD chimeras will also be instrumental in investigating functions of NC in budding, reverse transcription and integration^38,40,61^. Defining what enables specific chimeras to support or disrupt these steps will clarify essential NC activities for HIV-1 replication. Considering that a single dimeric gRNA in a virion is bound by a magnitude of viral (e.g. integrase and reverse transcriptase) and host RNA-binding proteins (e.g. APOBEC3 family), the ability to package gRNAs independent of NC by Gag-SRSF5 can help determine the interdependency of how these diverse RNA-binding proteins interact with the same gRNA. Given the potent dominant negative activities of Gag-RBD chimeras, understanding how they inhibit HIV-1 replication can reveal new vulnerabilities of HIV-1 for therapeutic development. Finally, our study also has implications for gene therapy vector design, as it raises the possibility that rational tuning of codon usage may improve packaging efficiency while maintaining functional protein output.

The unusual nucleotide composition of HIV-1 gRNA has, remarkably, remained stable through multiple decades and over 90 million infections of humans despite the HIV-1 genome being highly variable. Our study raises the possibility that selective packaging of the HIV-1 genome also contributes to maintaining this A-richness. Given HIV-1’s high mutation rate, a less restrictive mechanism of selecting viral genomes for packaging — through efficient virion assembly on viral RNAs with high adenosine bias rather than specific binding of viral proteins to a packaging signal made up of conserved RNA sequences — could provide Gag with a general mechanism for selective detection and packaging of genomes despite their constant diversification. Besides HIV-1, a large number of other viruses with RNA genomes (e.g. HTLV-1 and Rubella) also have a biased nucleotide composition^62^. While our study suggests that nucleotide composition may serve as a molecular signature for HIV-1 genome packaging, similar principles are likely applicable to a broad range of RNA viruses.

## ACKNOWLEDGEMENTS

We thank Dr. Wandy Beatty and the Molecular Microbiology Imaging Facility at the Washington University School of Medicine for their assistance with confocal microscopy and transmission electron microscopy experiments. We thank Evan Sherer for her assistance with single cell imaging studies. H.R.V. was supported by the National Science Foundation Graduate Research Fellowship under Grant No. DGE-1745038, DGE-2139839 and the National Institute of Allergy and Infectious Diseases of the National Institutes of Health under Award Number T32AI007172. This work was supported by NIH grants U54 AI170660 (A.T., S.B.K., N.S.), R01 AI179691 (S.B.K., N.S.) and R56 AI110221 (N.S.).

CLIP-seq and RNA-seq data were deposited to the Gene Expression Omnibus Database with accession numbers <u>GSE281422</u>, <u>GSE281423</u>, and <u>GSE281424</u>.

## MATERIALS AND METHODS

HIV-1 NL4-3-based proviral plasmids bearing heterologous RNA-binding domains from hnRNP and SRSF proteins were generated by conventional cloning. These clones were subsequently transfected into HEK293T and HeLa cells to study expression and assembly properties as well as gRNA packaging through the assays detailed in SI Appendix, Materials and Methods.

