## Supplemental text and Figures for "The biased adenosine-rich content of the HIV-1 genome serves as a molecular signature that facilitates efficient packaging"

### MATERIALS AND METHODS

*Plasmids and cloning.*

pNL4-3-derived plasmids containing three copies of a HA-tag in the stalk region of MA [1] and with (PR-) or without (PR+) an inactivating mutation in the protease (PR) gene were modified to insert the heterologous RNA-binding domains as follows. First, the NC coding sequence was replaced by a NotI restriction endonuclease site through conventional cloning using outward BssHII and SphI sites (see list below). Sequences coding for the first four and last three amino acids of NC were left intact to facilitate the proper proteolytic cleavage of the inserted heterologous RNA-binding domains. PCR amplicons containing individual or the complete set of RNA-binding domains from hnRNP and SR proteins were inserted into the Not I site (see list below). Chimeras bearing a subset of RRMs were also subcloned into a previously described two-color virus backbone [2] using SphI and SacII cut sites for microscopy analyses.

Deletions within the 5' leader were generated by overlap-extension PCR mutagenesis. The overlap PCR fragments were restriction digested and subsequently cloned into pNL4-3/MA-3xHA backbone via conventional cloning. ΔΨ5 was built on the ΔΨ1 backbone. ΔΨ3 and ΔΨ6 was built on the ΔΨ2 backbone. AatII/SphI fragment derived from these ΔΨ plasmids were then cloned into Gag-SRSF5 backbone. Gag-SRSF5 mutants (mGGA-1, mGGA-2, mCCG-1, mCCG-2, mGGG, β3'mut, SRSF4 linker, SRSF4 RRM1, and SRSF4 RRM2) were generated using overlap-extension PCR mutagenesis, with PCR amplicons being cloned into the NotI site via ligation or Gibson Assembly. To generate β3''β4mut, SRSF5 RRM1-2 sequence was PCR amplified using reverse primer with the intended nucleotide changes and the PCR amplicon was then cloned into the NotI site. mGGA-1 and mGGA-2 overlap PCR amplicons were generated with 2 fragments; mCCG-1 and mCCG-2 were generated with 4 fragments; mGGG was generated with 3 fragments.

Primer pairs used in cloning are listed in the below Supplementary Table 1:

| Primer | Sequence |
|---|---|
| **ΔNC-NotI-F** | ACTGAGCGGCCGCAATTTTAGGAACCAAAGAAAG |
| **ΔNC-NotI-R** | ACTGAGCGGCCGCCTCTCTCAGTACAATCTTTCATTTG |
| **BssHII-F** | ATGGCGCCCGAACAGGGACT |
| **AatII-F** | CCTGACGTCTAAGAAACCATTATTATC |
| **SphI-R** | GCCCTGCATGCACTGGATGCAATCTATCCCA |
| **hnRNP A1-RRM1-F** | ACTGAGCGGCCGCCAGCTGAGGAAGCTCTTCATTG |
| **hnRNP A1-RRM1-R** | ACTGAGCGGCCGCCTCTGGAGACAGCTCTCTTTGG |
| **hnRNP A1-RRM2-F** | ACTGAGCGGCCGCAAAAAGATATTTGTTGGTGGC |
| **hnRNP A1-RRM2-R** | ACTGAGCGGCCGCCCAGGGCTTTTCTAACTTCACAG |
| **hnRNP F-RRM1-F** | ACTGAGCGGCCGCGGCTTTGTGGTCAAGCTCCG |
| **hnRNP F-RRM1-R** | ACTGAGCGGCCGCCGGACTTGAACACCTCAATG |
| **hnRNP F-RRM2-F** | ACTGAGCGGCCGCATGGATTGGGTGTTGAAGC |
| **hnRNP F-RRM2-R** | ACTGAGCGGCCGCCCTGGCTGCTCTTAAACACCTC |
| **hnRNP F-RRM3-F** | ACTGAGCGGCCGCCACTGTGTCCACATGAGGG |
| **hnRNP F-RRM3-R** | ACTGAGCGGCCGCCTGAATTCAAGAAGAGTTCTAT |
| **hnRNP K-KH1-F** | ACTGAGCGGCCGCATGGTTGAATTACGCATTCTG |
| **hnRNP K-KH1-R** | ACTGAGCGGCCGCCGATTTTCTTCAGAATTTCTC |
| **hnRNP K-KH2-F** | ACTGAGCGGCCGCGACTGCGAGTTGAGGCTGTTG |
| **hnRNP K-KH2-R** | ACTGAGCGGCCGCCGATGATCTTTATGCACTCTAC |
| **hnRNP K-KH3-F** | ACTGAGCGGCCGCATTATTACTACACAAGTAAC |
| **hnRNP K-KH3-R** | ACTGAGCGGCCGCCCTGCAGCAAATACTGTGCAT |
| **SRSF1-RRM1-F** | ACTGAGCGGCCGCTGCCGCATCTACGTGGGTA |
| **SRSF1-RRM1-R** | ACTGAGCGGCCGCCGCTTCGAGGAAACTCCACCC |
| **SRSF1-RRM2-F** | ACTGAGCGGCCGCAACAGAGTGGTTGTCTCTGG |
| **SRSF1-RRM2-R** | ACTGAGCGGCCGCCATCAACTTTAACCCGGATGTA |

| | |
|---|---|
| **SRSF3-RRM1-F** | ACTGAGCGGCCGCTGTAAGGTTTATGTAGGCAATC |
| **SRSF3-RRM1-R** | ACTGAGCGGCCGCCACCATTCGACAGTTCCACTCT |
| **SRSF4-RRM1-F** | ACTGAGCGGCCGCCCGCGGGTGTACATCGGCC |
| **SRSF4-RRM1-R** | ACTGAGCGGCCGCCGCCGCGGGCATGCTCAACAA |
| **SRSF4-RRM2-F** | ACTGAGCGGCCGCTACAGACTTATTGTGGAGAA |
| **SRSF4-RRM2-R** | ACTGAGCGGCCGCCTGGCTTGTCTTCAACTAATCT |
| **SRSF5-RRM1-F** | ACTGAGCGGCCGCTGTCGGGTATTCATCGGGA |
| **SRSF5-RRM1-R** | ACTGAGCGGCCGCCAGCCCTAGCATGTTCAATAGT |
| **SRSF5-RRM2-F** | ACTGAGCGGCCGCAATCGTCTTATAGTTGAGA |
| **SRSF5-RRM2-R** | ACTGAGCGGCCGCCTTTGCTGCCTTCAATTAATTTTA |
| **Fragment 1-F (β3'mut, β3''β4mut, SRSF4 linker, SRSF4 RRM2, mGGA-1, mGGA-2, mCCG-1, mCCG-2, mGGG)** | TAATGATACAGAAAGGCAGCGGCCGCTGTCGGGTATTCATCGGG |
| **Fragment 1-R (β3'mut)** | GGTTCCATCAAGTTTTTCAATAGCATTCTTTAAGTCACCATAAG |
| **Fragment 2-F (β3'mut)** | TATTGAAAAACTTGATGGAACCGAAATAAATGGGAGAAAA |
| **Fragment 1-R (β3''β4mut)** | CCCTAAAAAATTAGCCTGGCGGCCGCCTGGCTTGTCTTCAATTAATTTTATTTTTC |
| **Fragment 1-R (SRSF4 linker)** | CGTAACTGCCATCTCGCCGTGGAGCCCTAGCATGTTCAATAG |
| **Fragment 2-F (SRSF4 linker)** | CCACGGCGAGATGGCAGTTACGGTTCTGGACGCAGTGGATATG |
| **Fragment 2-R (SRSF4 linker)** | AATTCTCAACTATAAGACGATTCTCTGTGCGAGTAGGAGGGC |
| **Fragment 3-F (SRSF4 linker)** | GCCCTCCTACTCGCACAGAGAATCGTCTTATAGTTGAGAATT |
| **Fragment 1-F (SRSF4 RRM1)** | TAATGATACAGAAAGGCAGCGGCCGCCCGCGGGTGTACATCGGC |

| | |
|---|---|
| **Fragment 1-R (SRSF4 RRM1)** | TACCTCTTCCACCTCGTGACCGGCCGCGGGCATGCTCAACAAT |
| **Fragment 2-F (SRSF4 RRM1)** | CGGTCACGAGGTGGAAGAGGTAGAGGACGATACTCTGA |
| **Fragment 1-R (SRSF4 RRM2)** | AATTCTCCACAATAAGTCTGTATTCTGTTCTTACAGGTGGAGC |
| **Fragment 2-F (SRSF4 RRM2)** | TGCTCCACCTGTAAGAACAGAATACAGACTTATTGTGGAGAATTTG |
| **Fragment 2-R (SRSF4 RRM2)** | CCCTAAAAAATTAGCCTGGCGGCCGCCTGGCTTGTCTTCAACTAATC |
| **Fragment 1-R (mGGA-1)** | GGCGGCGAGGGCCTGCCAGCTGACTCTTGAGGATAAATTCTC |
| **Fragment 2-F (mGGA-1)** | CAGCTGGCAGGCCCTCGCCGCCTTCATGAGACAAGCTGGGGAAG |
| **Fragment 1-R (mGGA-2)** | GTTCTGGAGGTTCTGCCAGCTGACTCTTGAGGATAAATTCTC |
| **Fragment 2-F (mGGA-2)** | CAGCTGGCAGAACCTCCAGAACTTCATGAGACAAGCTGGGGAAG |
| **Fragment 1-R (mCCG-1, mCCG-2)** | CTCTACAGGTGGAGCATTTCGTCTATCATTTCGAGGTCTG |
| **Fragment 2-F (mCCG-1, mCCG-2)** | ACGAAATGCTCCACCTGTAGAGAGAACAGAAAATCGTCTTATAG |
| **Fragment 2- R (mCCG-1)** | GGCGGCGAGCTTCTGCCAGCTGACTCTTGAGGATAAATTCTC |
| **Fragment 3- F (mCCG-1)** | CAGCTGGCAGAAGCTCGCCGCCTTCATGAGACAAGCTGGGGAAG |
| **Fragment 3-R (mCCG-1, mCCG-2) & Fragment 2-R (mGGG)** | CTCCTCCCCATTTATTTCCTTTCCAGAAAGTTTTTCAATAGC |
| **Fragment 4-F (mCCG-1, mCCG-2) & Fragment 3-F (mGGG)** | AAAGGAAATAAATGGGGAGGAGATAAAATTAATTGAAGGC |
| **Fragment 2-R (mCCG-2) &** | ATCTTTGAGCTTCTGCCAGCTGACTCTTGAGGATAAATTCTC |

| | |
|---|---|
| **Fragment 1-R(mGGG)** | |
| **Fragment 3-F (mCGG-2) & Fragment 2-F (mGGG)** | CAGCTGGCAGAAGCTCAAAGATTTCATGAGACAAGCTGGGGAAG |
| **Fragment 2-R (β3'mut, SRSF4 RRM1, mGGA-1, mGGA-2) & Fragment 4-R (mCCG-1, mCCG-2) & Fragment 3-R (mGGG, SRSF4 linker)** | CCCTAAAAAATTAGCCTGGCGGCCGCCTTTGCT |

**Supplemental Table 1:** Primer pairs for cloning Gag-RBD chimeras in this study.

*Cells, transfections, and viruses.*

HEK293T (ATCC CRL-11268), HeLa (ATCC CCL-2) and HeLa-derived TZM-bL (NIH AIDS Reagent Program) cells were maintained in Dulbecco's Modified Eagle Medium (DMEM) supplemented with 10% fetal bovine serum (FBS) at 37°C. MT4-GFP indicator cells containing an HIV-1 LTR-driven GFP reporter gene were described before[3] and grown in RPMI-10% FBS supplemented with puromycin. For virus production, HEK293T cells were transfected with the indicated proviral plasmid using polyethylenimine (PEI) and viral particles were harvested at 48 hours post transfection from filtered cell culture supernatants. HeLa cell transfections were performed with either PEI or Lipofectamine 3000 (Thermo Fisher Scientific) reagent according to the manufacturer's instructions.

*Immunoblotting.*

Cells were lysed in 1X radioimmunoprecipitation assay (RIPA) buffer and cleared lysates were diluted in 1X final sodium dodecyl sulfate (SDS) sample buffer. Viruses were pelleted on a 20% sucrose cushion prepared in 1X PBS and similarly resuspended in 1X sodium dodecyl sulfate

(SDS) sample buffer. Samples were separated by electrophoresis on Bolt 4–12% Bis-Tris Plus gels (Life Technologies), transferred to nitrocellulose membranes and probed with the following antibodies in Odyssey/Intercept Blocking Buffer (LI-COR): mouse monoclonal anti-CA (183-H12-5C, NIH AIDS reagents), mouse monoclonal anti-HA (Biolegend, clone 16B12), goat polyclonal anti-Env (ARP, 12-6205-1), rabbit polyclonal anti-Nef (NIH AIDS Reagents, ARP2949), mouse monoclonal anti-β-tubulin (Santa Cruz, SC-5274), mouse monoclonal anti-actin (Santa Cruz, SC-8432), mouse monoclonal anti-$Na^+$-$K^+$ ATPase (Santa Cruz, SC-21712). Membranes were probed with fluorophore-conjugated secondary antibodies and visualized using the LI-COR Odyssey system. Where indicated, Gag protein in cells and virions were quantified using the LI-COR Image Studio software to normalize for CLIP protein:RNA adduct signal.

*RNA extraction and RT-qPCR.*

Cell and virion associated RNA was harvested with TRIzol (Thermo Fisher Scientific) and precipitated overnight in isopropanol. RNA was resuspended and reverse transcribed with the High-Capacity cDNA Reverse Transcription Kit (Applied Biosystems) according to the manufacturer's instructions. qPCR was performed using PowerUp 2X SYBR Green Master Mix (Applied Biosystems). Following primers in Supplemental Table 2 were used to quantify their respective targets:

| Primer | Sequence |
|---|---|
| **Unspliced/Gag-Pol-F** | TTCTTCAGAGCAGACCAGAGC |
| **Unspliced/Gag-Pol-R** | GCTGCCAAAGAGTGATCTGA |
| **Multiply spliced-F** | TCTATCAAAGCAACCCACCTC |
| **Multiply spliced-R** | CGTCCCAGATAAGTGCTAAGG |
| **7SL RNA-F** | CACCAGGTTGCCTAAGGAGGGG |
| **7SL RNA-R** | TCCCACTACTGATCAGCACGGG |

| | |
|---|---|
| **GAPDH-F** | AGGTGAAGGTCGGAGTCAACG |
| **GAPDH-R** | GGTCATTGATGGCAACAATATCCACTTTAC |
| **RPL36A-F** | GCACCAACCCCATAAAGTGAC |
| **RPL36A-R** | CTGGGCGTACAGAGAATCCT |
| **RPS6-F** | TGGACGATGAACGCAAACTTC |
| **RPS6-R** | TTCGGACCACATAACCCTTCC |
| **RPS16-F** | TCGGACGCAAGAAGACAGC |
| **RPS16-R** | AGCAGCTTGTACTGTAGCGTG |

**Supplemental Table 2:** RT-qPCR primers used in this study.

*Quantification of reverse transcription and integration*

MT4-GFP cells were grown in 24-well plates and infected with VSV-G pseudotyped WT or Gag chimera viruses at an MOI of 0.5 or 1 in the presence of polybrene. Cells were collected at 24 hpi, washed with PBS, and resuspended in 200 µL lysis buffer (100 mM NaCl, 10 mM Tris-HCl pH 8, 25 mM EDTA pH 8, 0.5% SDS, 0.1 mg/mL Proteinase K). Samples were incubated at 50°C by constant agitation (2,000 rpm) in a thermal mixer (Eppendorf) for 2 h. DNA was extracted by phenol-chloroform extraction. Extracted DNA was analyzed for accumulation of RT products, 2-LTR circles, and integration as described before [4,5].

*Northern blotting*

For northern blotting experiments, 293T cells were transiently transfected with 5 μg of each plasmid using a 4:1 ratio of PEI and 150 mM NaCl. Media was replaced 24- and 44-hours post-infection. 4 hours after the 44-hour media change, extracellular viral particles were harvested by passing culture supernatants through a 0.22 um filter. Particles were then concentrated via ultracentrifugation through a 20% sucrose cushion. RNA was isolated from transfected cells and concentrated virions using Trizol Reagent. Northern blot analysis was performed using a modified version of a previously described protocol. Briefly, RNA samples were incubated in 1x MOPS electrophoresis buffer, 8% formaldehyde, 50% formamide, and gel loading buffer and then heated at 85°C for 10 minutes. The denatured samples were loaded into the wells of a 1% agarose gel containing 2.2M formaldehyde, and electrophoresis proceeded for approximately 7 hours at 50V. RNA species were then vertically transferred to a nylon membrane. Following transfer, the membrane was exposed to UV to induce crosslinking. The membrane was then incubated in prehybridization buffer (50 mM Tris, pH 7.5, 120 μg/mL salmon sperm DNA) at 55°C. After 4 hours, a $^{32}$P-labelled probe complementary to the viral genome or 7SL RNA was added to the prehybridization buffer and allowed to hybridize with the membrane at 55°C overnight. The membrane was washed the next day and then exposed overnight in a phosphorimaging cassette. The screen was imaged using an Amersham™ Typhoon™ scanner.

*Membrane Flotation Assays.*

Membrane flotation assays were performed as previously described [6]. HEK293T cells grown in 10 cm plates were transfected with 10μg of the indicated proviral plasmids using PEI and cell culture media was replaced 6-8 hours post-transfection. 24 hours post transfection, cells were collected, washed once with 1X STE buffer (100 mM NaCl, 10 mM Tris-Cl pH 8.0, 1 mM EDTA). Afterwards, cells were resuspended in 500μl of hypotonic solution (10mM Tris [pH 7.4], 10mM KCl, 1 mM EDTA) supplemented with Complete Protease Inhibitor Cocktail (Roche), and placed on ice for 15 minutes. Cells were then lysed by dounce homogenization and centrifuged at 1000 x g for 5 minutes, 4°C. 350μl of post-nuclear supernatant was collected and mixed with 1.65 mL of 90% sucrose solution. 6.5 mL of 65% sucrose solution followed by 2.5 mL of 10% sucrose solution were layered on top of the mixture. Centrifugation was then performed with a Beckman SW41 Ti rotor at 35,000 rpm for 4-18 hours, 4°C. 1 mL fractions were collected, with the 3rd fraction and 11th fraction from top corresponding to the membrane and cytosol, respectively. 100 μL of each fraction was kept for RNA analysis while the remaining 900 μL was used for protein precipitation. Aliquots for RNA analysis were treated with Proteinase K for 10 min at 37°C and RNA was extracted with an equal volume of phenol:chloroform:isoamyl alcohol (125:24:1) (Sigma-Aldrich) and precipitated with the addition of 1 µL Glycoblue (Ambion), 0.25 volume of 3M NaOAc pH 5.2, and 2.5 volumes of 1:1 ethanol:isopropanol. The resulting RNA samples were treated with DNase, reextracted using one more round of phenol:chloroform extraction and used for RT-qPCR. Protein was precipitated in 10% trichloroacetic acid (TCA) for 2-4 hours and pelleted by centrifugation. Protein pellets were washed twice with 10% TCA and once with ice cold acetone before being resuspended in SDS-PAGE sample buffer and used for western blot analyses.

*Equilibrium density sedimentation of virion core components in vitro*

Equilibrium density sedimentation of virion core components was performed as previously described [7]. Briefly, HEK293T cells grown on 10-cm dishes were transfected with the above-mentioned plasmids. 48 hours post-transfection, cell-free virions collected from cell culture supernatants were pelleted through a 20% sucrose cushion. Pelleted virions were resuspended in 1X PBS and treated with 0.5% Triton X-100 for 2 minutes at room temperature. Immediately after, samples were layered on top of a 30-70% linear sucrose gradients in 1X STE buffer and centrifuged for 16 hours at 4°C and 28,500 rpm using an SW55Ti rotor. 500 µL fractions collected from the top were analyzed by immunoblotting using a mouse monoclonal anti-HIV p24 antibody (183-H12-5C, NIH AIDS reagents) and Q-PCR-based assays for RT activity [8].

*Transmission electron microscopy.*

To visualize virus assembly sites in cells, HEK293T cells were grown in 10 cm plates and transfected with 10µg of proviral plasmid as previously described. 24 h post-transfection, cells were harvested and fixed in 2% paraformaldehyde/2.5% glutaraldehyde (Ted Pella Inc., Redding, CA) in 100mM cacodylate buffer, pH 7.2 for 2 h at room temperature. Samples were washed in cacodylate buffer and postfixed in 1% osmium tetroxide (Ted Pella Inc.)/1.5% potassium ferricyanide (Sigma, St. Louis, MO) for 1 h. Samples were then rinsed extensively in dH20 prior to en bloc staining with 1% aqueous uranyl acetate (Ted Pella Inc.) for 1 h. Following several rinses in water, samples were dehydrated in a graded series of ethanol and embedded in Eponate 12 resin (Ted Pella Inc.). Ultrathin sections of 95nm were cut with a Leica Ultracut UCT ultramicrotome (Leica Microsystems Inc., Bannockburn, IL), stained with uranyl acetate and lead citrate. To visualize viral particles, HEK293T cells grown in 15-cm plates were transfected with 30 µg of the indicated plasmid. Cell free virions were harvested from the supernatant 48 h post transfection, filtered through 0.22 µm filters and pelleted through a 20% sucrose cushion via centrifugation using a Beckman SW32-Ti rotor at 28,000 rpm for 1.5 h, 4°C. Virion pellets were fixed and processed for EM as described above. All samples were viewed on a JEOL 1200 EX transmission electron microscope (JEOL USA Inc., Peabody, MA) equipped with an AMT 8-megapixel digital camera and AMT Image Capture Engine V602 software (Advanced Microscopy Techniques, Woburn, MA).

*Immunofluorescence microscopy.*

HeLa cells were plated in 24-well glass-bottom dishes (Mattek Corporation, Ashland, MA, USA) coated with poly-L-lysine (50,000 cells/well) and transfected with 500ng of the indicated proviral plasmid using either PEI or Lipofectamine 3000 transfection reagent. At 24 h post-transfection cells were fixed with 4% paraformaldehyde in PBS, for 10 min and, for immunofluorescence-based experiments, permeabilized with 0.1% Triton X-100 in PBS for 10 minutes. Permeabilized cells were blocked with 5% goat serum in PBS-T (PBS, 0.1% Tween 20) blocking solution for 30 min at room temperature. Cells were probed with a mouse monoclonal anti-HA (Biolegend, clone 16B12) in blocking solution for 1-2 h at room temperature, followed by goat anti-mouse Alexa Fluor 594 secondary antibody (1:1000, Thermo Fisher Scientific) for 1 h at room temperature.

Nuclei were stained with DAPI for 10 min. Cells were visualized with a Zeiss LSM880 Confocal Microscope with Airyscan, oil immersion x63 or A1R Confocal Microscope, oil immersion x60.

*Single virion analysis*

To generate labeled, two-colored virus-like particles for single virion analysis, approximately 500,000 HEK293T cells were plated in each well of a six-well dish and transfected with plasmids encoding wild-type, SRSF5 RBD, or ΔNC two-color self-tagging viruses using PEI. The media was exchanged at 24 h post-transfection. Virus particle-containing supernatants were harvested at 48 h post-transfection, filtered through a 0.22-μm filter, and centrifuged through 20% sucrose for 2 h at 18,213 rpm. The medium was discarded after centrifugation, and concentrated viral particles were resuspended in 1× PBS, plated in a 24-well glass-bottom dish (Cellvis, Mountain View, CA), and left overnight at 4°C to allow virus particles to settle down on the glass wells. Microscopy was performed using a Nikon Ti-Eclipse inverted wide-field microscope (Nikon Corp, Minato, Tokyo, Japan) using a 100× Plan Apo oil objective lens (numerical aperture [NA] 1.45). Cell and virion images were captured using an ORCA—Flash4.0 CMOS camera (Hamamatsu Photonics, Skokie, IL, USA) and the following excitation/emission filter sets: 510/535 nm (YFP) and 585/610 nm (mCherry). All images were processed and analyzed using FIJI/ImageJ2. FIJI/ImageJ2's built-in tools were used to measure the mean fluorescent intensities (MFIs) of both the virions and the background by selecting regions of interest in the Gag-YFP channel and the corresponding areas in the MS2–mCherry (unspliced RNA) channel. Virion background subtracted MFIs for MS2–mCherry and Gag–YFP channels were plotted using GraphPad Prism (version 10.1.2). The data were plotted for MS2–mCherry MFI, Gag–YFP MFI, and MS2–mCherry/Gag–YFP MFI.

*RNA-seq.*

HEK293T cells were grown in 24-well plates and transfected with 400ng of the indicated proviral plasmid using PEI. 48 h post transfection, cell culture media was collected and filtered through 0.22 μm filters, before being pelleted through a 20% sucrose cushion at 15,000 rpm for 1.5 h, 4°C. RNA was harvested from purified cell free virions with TRIzol as previously described. RNA samples were prepared for sequencing using the TruSeq Stranded Total RNA Sample Prep Kit (Illumina) according to the manufacturer's instructions but without rRNA depletion and as detailed previously [9]. Barcoded samples were pooled and sequenced using an Illumina NextSeq platform (1x75) at the Center for Genome Sciences at Washington University.

*Photoactivatable ribonucleoside-enhanced cross-linking and immunoprecipitation (PAR-CLIP).*

PAR-CLIP experiments were conducted as previously described with some modifications [1,10,11]. HEK293T cells were grown in 10-cm plates and transfected with 10 μg of the indicated proviral plasmid. Cell culture media was replaced 6-8 h post-transfection, and 4-thiouridine (4SU) was added to the cell culture medium at a concentration of 100 μM 16 h before harvesting cells and virions. 48 hours post-transfection, virions and cells were collected. Virions in the cell culture supernatant were filtered through 0.22 μm filters and pelleted through a 20% sucrose cushion at 28,000 rpm for 1.5 hr, 4°C. Transfected cells were harvested with a cell scraper. Both cells and purified virions were resuspended in PBS and protein-RNA complexes UV-crosslinked (365nm) before being lysed in 1X RIPA lysis buffer. Cell and virus lysates were then treated with RNase A (Thermo Fisher Scientific) and DNase I (Roche) at a final concentration of 20and 60 U/mL, respectively. HA-tagged Gag proteins were immunoprecipitated using mouse monoclonal anti-HA antibody conjugated Protein G-coupled Dynabeads (Thermo Fisher Scientific), as previously described. Gag associated RNAs were radiolabeled with γ-$^{32}$P-ATP using T4 Polynucleotide Kinase (New England Biolabs). The protein-RNA adducts were separated via SDS-PAGE and transferred onto nitrocellulose membranes before being visualized with autoradiography film. Protein-RNA adducts were excised from nitrocellulose membranes and digested with Proteinase K, and RNA was extracted with phenol:chloroform:isoamyl alcohol (125:24:1) (Sigma-Aldrich). Resultant RNA was pelleted with the addition of 1µL Glycoblue (Ambion), 0.25 volume of 3M NaOAc pH 5.2, and 2.5 volumes of 1:1 ethanol:isopropanol. The RNA pellet was washed with 80% ethanol, dried and dissolved in 13 µL RNase-free H2O. RNA was ligated with barcoded adapters and ligation products were separated and purified on Novex 15% TBE-Urea gel (Invitrogen) as previously described with some exceptions [10]. Here we used barcoded 3' adapters and thus samples were pooled into groups of two prior to 5' adapter ligation (Supplementary Table 3). Furthermore, the amount of 3' adapter added to each reaction was reduced to 5pmole. An end-labeled 15-nucleotide RNA oligonucleotide was prepared in parallel as positive control for adapter ligations. The resulting ligation products were reverse transcribed using Superscript III First Strand Synthesis Kit (Invitrogen), and cDNA was amplified using Phusion Polymerase (New England Biolabs). PCR products were gel purified and sequenced on a NextSeq platform (1x75bp) at the Center for Genome Sciences at Washington University.

| **Oligos** | **Sequence** |
|---|---|
| CLIP1 | 5′adenylated/NNTGACTG TGG AAT TCT CGG GTG CCA AGG-3′dideoxyC |
| CLIP2 | 5′adenylated/NNACACTCTGG AAT TCT CGG GTG CCA AGG-3′dideoxyC |
| CLIP3 | 5′adenylated/NNACAGAG TGG AAT TCT CGG GTG CCA AGG-3′dideoxyC |
| CLIP4 | 5′adenylated/NNGCGATA TGG AAT TCT CGG GTG CCA AGG-3′dideoxyC |
| CLIP31 | 5′adenylated/NNTAGCGA TGG AAT TCT CGG GTG CCA AGG-3′dideoxyC |
| CLIP32 | 5′adenylated/NNCTGTAG TGG AAT TCT CGG GTG CCA AGG-3′dideoxyC |
| CLIP33 | 5′adenylated/NNTAGTCG TGG AAT TCT CGG GTG CCA AGG-3′dideoxyC |
| CLIP34 | 5′adenylated/NNAGTGTC TGG AAT TCT CGG GTG CCA AGG-3′dideoxyC |
| CLIP35 | 5′adenylated/NNATCGAC TGG AAT TCT CGG GTG CCA AGG-3′dideoxyC |

| | |
|---|---|
| CLIP36 | 5′adenylated/NNGACATG TGG AAT TCT CGG GTG CCA AGG-3′dideoxyC |
| CLIP37 | 5′adenylated/NNGACTAC TGG AAT TCT CGG GTG CCA AGG-3′dideoxyC |
| CLIP38 | 5′adenylated/NNATCTCG TGG AAT TCT CGG GTG CCA AGG-3′dideoxyC |
| 5’ adapter | rGUU CAG AGU UCU ACA GUC CGA CGA UC |
| RT primer | GCCTTGGCACCCGAGAATTCCA |
| cDNA PCR (F) | AATGATACGGCGACCACCGAGATCTACACGTTCAGAGTTCTACAGTCCGA |
| cDNA PCR (R) | CAAGCAGAAGACGGCATACGAGATCGTGATGTGACTGGAGTTCCTTGGCACCCGAGAATTCCA |
| Control oligo | rAUAGCUACGAUUGCA |

**Supplemental Table 3:** Sequences of adapters and primers used in CLIP-seq assays.

*Bioinformatic analysis of NGS data*

**Clip-seq:**

Mapping:

Adapter trimming was performed using BBDuk from BBTools version 39.06. The 3′ adapter sequences were removed with parameters k=8 and mink=7 to define the k-mer size and minimum k-mer size. Additional parameters included ml=18 and maxlength=52 to filter reads based on length. The rcomp function was disabled to only consider kmers in the forward orientation.

Trimmed reads were further processed with BBDuk for separation based on their 3′ barcode sequences. Parameters included a k-mer size of 8, maskmiddle=f to ensure only exact matches were trimmed, and copyundefined was set to handle undefined bases. The resulting FASTQ files are the trimmed reads specific to each barcode.

Next, the trimmed reads were mapped to their corresponding Gag chimera viral genomes and WT viral genome (HIV-1$_{NL4-3}$/PR$^{-}$) with Bowtie version 1.3.1. Uniquely mapped reads were reported, allowing one mismatch (-v 1 -m 1). Mapped reads were processed with the mpileup function form SAMtools to generate pileups for HIV-$1_{NL4-3}$/$PR^{-}$, and these pileups were then analyzed using customized in-house scripts and visualized as viral density maps with GraphPad Prism (v6). Reads that failed to map to their corresponding viral genomes were extracted using SAMtools (-b -f 4) and then aligned to the hg19 human genome with Bowtie, allowing up to 10 multi-mapped reads with one mismatch. Various file formats of mapped reads were generated, including BED and SAM files, for further analysis.

Cluster Generation by PARalyzer and Cluster Annotations

PARalyzer was used for peak calling using SAM or BOWTIE outputs to generate clusters, which represent binding sites derived from overlapping mapped reads. Clusters were annotated using publicly available databases, ENSEMBL v72 and tRNASCANse [12].

For a cluster or a read that mapped to a location with multiple gene features (i.e., intron-exon or coding-sequence-3′ UTR junctions), we required greater than half the length of the read/cluster to map to the assigned class. In other cases where a cluster/read could be assigned to multiple gene features due to the presence of alternatively spliced isoforms in the ENSEMBL database, the following order of preference was chosen: coding sequence, 5′UTR, 3′UTR, intron. In some experiments, the number of reads associated with each cluster was taken as a measurement of how frequently a given cluster (i.e., binding site) was bound by Gag. To identify sequence motifs within mRNAs, PARalyzer parameters were adjusted to generate clusters with a mean length of ~10 nucleotides surrounding the crosslinking site and which overlapped > 95% with the clusters generated above. Perl scripts were written for comparison of clusters in two separate data sets, determining the nucleotide composition of clusters and identifying features of mRNA molecules (i.e., 5′UTR, 3′UTR, CDS length) bound by Gag.

Correlation Analysis

Cross-correlation analysis was performed to measure the similarity of Gag binding frequencies between two datasets (Oppenheim & Schafer, 1975). The correlation function (CF(s)) compared the signals by shifting one dataset relative to the other and calculating their normalized correlation. The search window for correlation analysis was set to 400 nucleotides.

To assess the statistical significance of observed correlations, Monte Carlo simulations were used to generate 10,000 randomized datasets with the same structure as the original (i.e., identical numbers of binding sites and reads per site). Confidence intervals were calculated for p-values of 0.05 based on the cross-correlation results of these randomized samples.

Motif analysis

The bed files were annotated and reads in the mRNA regions were extracted using in-house scripts. These extracted reads were then converted to FASTA format using BEDTools for the subsequent analysis.

Motif analysis was conducted using MEME Suite with the following parameters: -nmotifs 10 -minw 5 -maxw 20 -dna. The parameters specified the identification of up to 10 motifs with a minimum width of 5 and a maximum width of 20 base pairs, applied to DNA sequences.

**RNA-seq:**

Total RNA-seq reads were first mapped to the HIV-1 genome using Bowtie. For the remaining non-viral reads (which we considered here as 100% to better depict the cellular RNA species present in each sample) they were mapped to ribosomal cellular RNA **(rRNA, Fig. S1F)**, non-rRNA cellular RNA **(Fig. 1H)** or constituted of reads that remained unmapped. Lastly, the non-rRNA cellular reads were further classified as 7SL RNA, mRNA, or Other **(Fig. S1G)**. The STAR aligner was used to map the RNA-seq data to human genome indices (hg19) with a mismatch rate threshold of 4%. Mapped reads were than annotated with featureCounts.

All CLIP-seq and RNA-seq data were analyzed using publicly available programs (BBDuk toolkit, Bowtie, and SAMtools) and in-house scripts [1,10]. Briefly, reads that contained ambiguous nucleotides, did not contain the 3′ adapter sequence or were shorter than 10nt were discarded. The remaining reads were then separated based on their 3′ barcode sequences and collapsed to generate a set of unique sequences. Reads were then aligned to the human genome (hg19) or to corresponding viral genomes, with up to 1 mismatch allowed using the Bowtie algorithm [13]. In the case of reads that mapped to the human genome, locations were only reported for those with the minimum number of observed mismatches for each read (Bowtie criteria: -v 1 -m 10 for mapping to hg19, and - v 1 -m 1 for mapping to the viral genomes). SAMtools was used to generate pileups of mapped reads, which were subsequently processed using in-house scripts and visualized by Prism GraphPad software (v6) [14].

### SUPPLEMENTAL FIGURE LEGENDS

**Figure S1. Properties of Gag-RBD chimeras with heterologous RNA binding modules in place of NC. A.** HEK293T cells were transfected with HIV-$1_{NL4-3}$ proviral plasmids encoding the indicated Gag-RBD chimeras. Cell lysates were analyzed with immunoblotting for Nef and Env (gp120). **B.** The indicated HIV-1 particles bearing Gag-RBD chimeras were titered on TZM-bl reporter cells, with 100 μg/mL dextran-sulfate added at 6-16 hpi to limit infection to a single cycle. The titers were subsequently normalized based on particle numbers as quantified by a reverse transcriptase (RT) activity assay and presented relative WT (set to 1). Titers are mean ± SEM from n=3 independent biological replicates. **C.** Densitometry analysis of Gag:RNA adducts from PAR-CLIP autoradiograms from Fig. 1B. Data are normalized relative to immunoprecipitated protein levels and graphed as fold difference relative to WT (set to 1). Graph shows the average of 3-5 independent experiments, error bars show the SEM. **D, E.** RT-qPCR quantification of relative copies of gRNA, multiply spliced viral RNAs, and 7SL RNA (7SL) from purified virions without (D) or with (E) normalization relative to virus particle numbers quantified based on RT activity. The columns represent the average of 3 independent experiments, error bars represent SEM. **F.** Classification of non-rRNA cellular RNA reads from virion total RNA-seq experiments as 7SL, mRNA, and Other. **G.** Northern blot analysis of unspliced (US), partially spliced (PS), multiply spliced (MS) HIV-1 RNAs and host 7SL RNA isolated from transfected cells and purified HIV-1 particles. **H-J.** The indicated virions were subjected to equilibrium density sedimentation following treatment with 0.5% Triton as detailed in Materials and Methods. Ten fractions harvested from the top of sucrose gradients were analyzed via immunoblotting using a mouse monoclonal anti-CA antibody (H, I) or a qPCR assay for RT activity (J) or CA content via (F). Results show the mean values ± SEM of three independent biological replicates. **K.** MT4-GFP cells were infected with HIV-1/VSV-G at MOI of 1 in the presence and absence of nevirapine (NVP) at 25 μM. At 24 hpi, cells were collected for DNA extraction as detailed in Materials and Methods. Extracted DNA was analyzed by qPCR for accumulation of early RT products (ERT), late RT products (LRT), 2-LTR circles (2-LTR), and integrated viral DNA (Alu-PCR). Data show the mean from two independent replicates. **P*<0.05; ** *P*<0.01; **** *P*<0.0001 by one-way ANOVA with Dunnet's multiple comparison test for correction.

**Figure S2. Trafficking and assembly properties of Gag-RBD chimeras. A.** Representative (n=3) immunoblot analysis of $Na^{+}K^{+}$ ATPase and β-tubulin from membrane (M) and cytoplasmic (C) fractions isolated from 293T cells transfected with the indicated Gag-RBD proviral plasmids (PR null). Samples are the same as those shown in Fig. 2A. **B.** Representative confocal microscopy images of HeLa cells transfected with proviral plasmids encoding the indicated Gag-RBD. Cells were stained for Gag (red) and nuclei by DAPI (blue). **C.** Representative fluorescence microscopy images of HeLa cells transfected with dually labeled proviral constructs. Scale bars represent 20 μm. ROIs are indicated by white boxes. **D.** Confocal imaging of the bottom surface of cells transfected with two-color Gag-YFP-SRSF5/MS2-mCherry or Gag-YFP-hnRNPA1/MS2-mCherry viruses.

**Figure S3. RNA binding properties of Gag-SRSF5 gRNA in cells. A.** Proportion of reads that map to cellular and viral genomes obtained from three independent Gag-CLIP experiments performed on cell lysates. Each point represents an independent experiment. **B.** Representative sequence motifs most frequently occurring in Gag-bound cellular mRNA clusters generated from CLIP experiments. **C.** Read density distribution on full-length HIV-1 gRNA (x-axis) from CLIP-seq experiments in which WT Gag and Gag-RBD chimeras (PR null backbone) were immunoprecipitated from transfected 293T cell lysates. **D.** Depiction of secondary structures within the 5'-UTR of HIV-1 RNA (top) and table (bottom) showing sequences deleted in ΔΨ1-6 mutants used for this study. TAR, trans-activation response element; PBS, primer-binding site; DIS, dimerization initiation signal; SD, major splice donor. **E.** RT-qPCR analysis of relative copies of virion-incorporated HIV-1 gRNA. The gRNA levels of ΔΨ mutants derived from WT Gag and Gag-SRSF5 backbones are graphed as fold differences relative to Ψ/WT Gag and Ψ/Gag-SRSF5, respectively (both set as 1). The columns represent the average of two independent experiments (each done in one to two technical replicates representing independent transfections), and the error bars represent SEM. *$P<0.05$; **$P<0.01$; ***$P<0.001$; by one-way ANOVA with Dunnet's multiple comparison test for correction. Only the significant comparisons are shown. **F.** Representative immmunoblot analysis of purified virus particles (PR null backbone) using a mouse monoclonal anti-HA antibody.

**Figure S4. Analysis of the RNA binding properties of Gag-RBD chimeras at the PM and immature virions. A-B.** Densitometry analysis of Gag:RNA adducts from autoradiograms (See Figure 3C, 3D) of independent CLIP replicates done on PM fractions (A) or immature virus particles (B). Data are normalized relative to immunoprecipitated protein levels and graphed as fold differences relative to WT (set to 1). Graph shows the average of 3 independent experiments and the error bars represent SEM. **C.** Proportion of reads that map to cellular and viral genomes obtained from CLIP experiments done on virus lysates. Each point represents an independent experiment. **D.** Correlation analysis of WT, ΔNC-NC, and Gag-SRSF5 binding to viral RNA in virions from independent CLIP experiments. **E.** Classification of read clusters (top) and total number of reads in each cluster (bottom) generated from virion Gag-CLIP experiments that map to cellular RNAs; total number of reads is indicated below each pie chart**. F.** Representative sequence motifs most frequently occurring in Gag-bound cellular mRNA clusters generated from virion Gag-CLIP experiments. **G.** Frequency distribution of nucleotide occurrence (read density) in reads mapping to the HIV-1 genome derived from CLIP experiments done on PM fractions.

**Figure S5. Rational design of Gag-SRSF5 variants with reduced affinity for GGA and enhanced affinity for CCG or GGG. A.** Structural depiction of key residues involved in GGA motif recognition in Gag-SRSF5 variants with reduced GGA affinity (mGGA1- and mGGA-2) or enhanced affinity for CCG or GGG motif (mCCG-1, mCCG-2, and mGGG) are shown. The amino acid positions shown are based on full-length SRSF5. **B-E.** Representative autoradiograms of Gag-RNA complexes (top) and immunoblot analysis of Gag-SRSF5 mutants (bottom) immunoprecipitated from cells expressing the indicated Gag chimeras (B) and progeny virus particles (D). (*) marks the IgG-heavy chain from the immunoprecipitating antibody. Eluates from IPs were normalized to load similar amounts of protein for Gag and Gag-SRSF5 variants to more clearly illustrate RNA binding ability per protein. Densitometry analysis of Gag:RNA adducts from autoradiograms (e.g. panels B and D) of independent replicates. Data are normalized relative to immunoprecipitated protein levels and graphed as fold differences relative to WT (set to 1). Graph shows the average of 3 technical replicates, representative of two independent experiments, and the error bars represent SEM. **F.** RT-qPCR quantification of relative copies of RPL36A, RPS6, and RPS16 RNA from HIV-1 particles collected from 293T cells expressing the indicated Gag chimeras. Data show the average of 3 independent experiments and the error bars represent SEM. Indicated statistical analyses were done by one-way ANOVA with Dunnet's multiple comparison test for correction, whereby n.s. denotes non-significant comparisons. Other group comparisons are not displayed due to clarity.

**Figure S6. Dominant negative activities of Gag-RBD chimeras. A.** Proviral plasmids encoding ΔGag or WT Gag was co-transfected with the indicated Gag-RBD proviral plasmids in 293T cells. 2 days post-transfection, cell lysates and purified virions were harvested for immunoblot analysis of Gag using a mouse monoclonal antibody against the HA-tag. Quantitation of virion immunoblot results from Fig. 5A is shown. Fold increase of the virion-incorporated Gag-RBD chimeric proteins in the presence of WT Gag over ΔGag co-transfection control was graphed. Data show the mean fold increase from n=2 independent experiments. **B-C.** 293T cells were transfected with equal ratio of Gag-RBD chimeric HIV-1 proviral plasmids and a WT Gag HIV-1 proviral plasmid. Particle release was quantified by and RT activity assay (B). RT-qPCR quantitation of relative HIV-1 gRNA and multiply spliced HIV-1 RNA copies from purified virions (C). Data show the mean from four independent replicates. *$P<0.05$, **$P<0.01$ by one-way ANOVA with Dunnet's correction. Non-significant comparisons are not displayed for clarity. **D.** 293T cells were co-transfected with a WT Gag proviral plasmid and variable amounts of the Gag-RBD chimeras. Particle release of samples shown in Fig. 5C was quantified by an RT activity assay.

**Figure S7. Gag-RBD chimeras bearing single RRMs bound RNA in cells, but do not package gRNA efficiently. A.** 293T cells were transfected with $PR^-$ pNL4-3 proviral plasmids encoding Gag chimeras in which the NC domain was replaced by single RRMs (e.g. M1, M2) from the indicated hnRNP and SRSF proteins. Cell lysates and purified virus particles were harvested 2 days post-transfection and analyzed by immunoblotting for Gag and Gag-Pol (GP). Gag-RBD:RNA complexes were immunoprecipitated from cell lysates and analyzed using autoradiography and immunoblotting. **B.** The relative titers of indicated Gag-RBD particles were determined via single-cycle infection of TZM-bl reporter cells. **C.** RT-qPCR quantification of relative copies of gRNA, multiply spliced HIV-1 RNA, and 7SL RNA from purified virions. **D-E.** 293T cells were transfected with equal ratio of Gag-RBD chimeric HIV-1 proviral plasmids and a WT Gag HIV-1 proviral plasmid. Particle release was quantified by and RT activity assay (D). RT-qPCR quantitation of relative HIV-1 gRNA and multiply spliced HIV-1 RNA copies from purified virions (E). **F.** 293T cells were co-transfected with a WT Gag proviral plasmid and variable amounts of the Gag-RBD chimeras. Particle release of samples shown in Fig. 5E was quantified by an RT activity assay. Data show the mean from four independent replicates. *$P<0.05$; **$P<0.01$; ****$P<0.0001$ by one-way ANOVA with Dunnet's correction. Non-significant comparisons are not displayed for clarity.

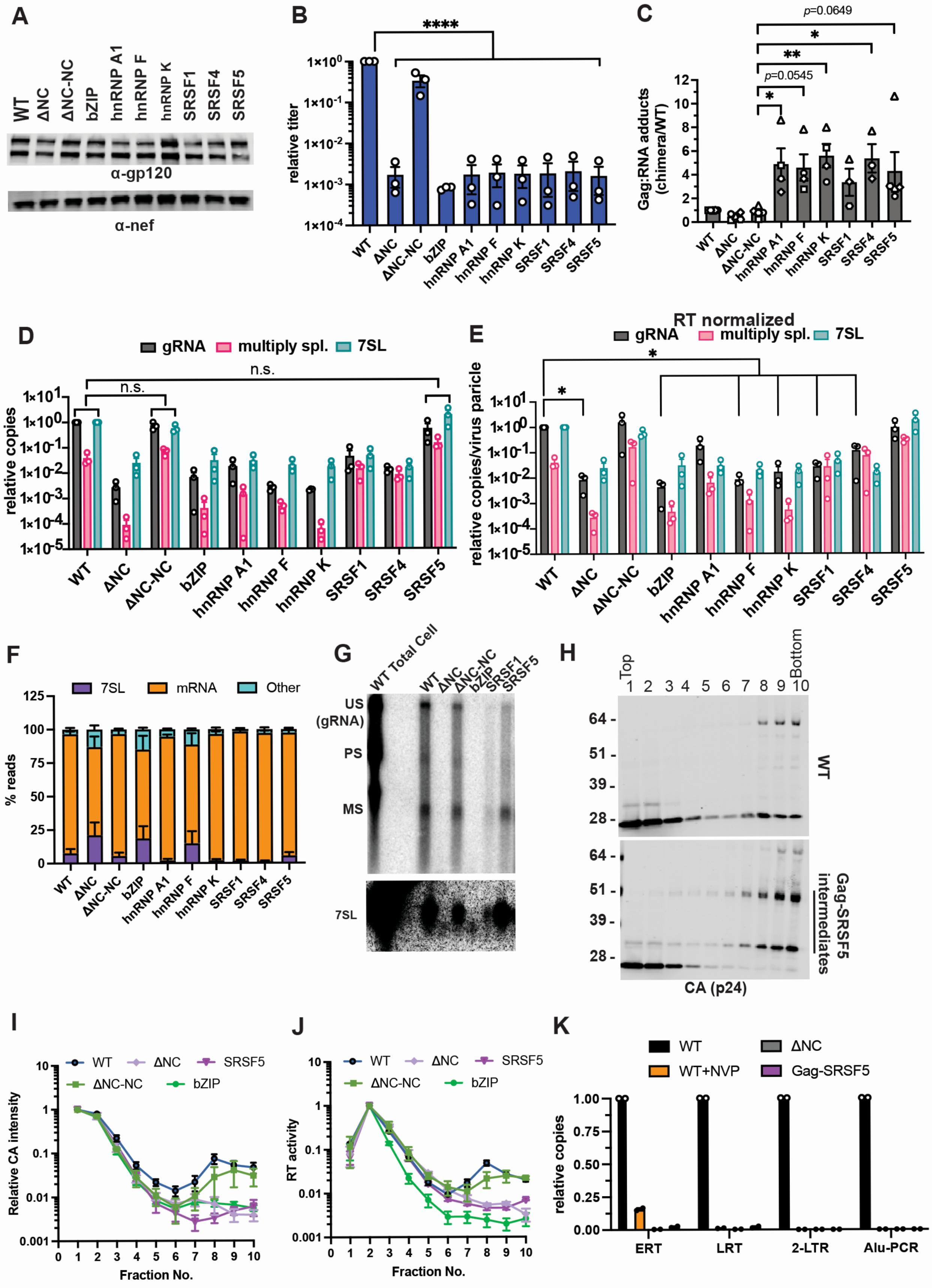

A
WT
ΔNC
ΔNC-NC
bZIP
hnRNP A1
hnRNP F
hnRNP K
SRSF1
SRSF4
SRSF5
α-gp120
α-nef
B
****
relative titer
C
p=0.0649
*
**
p=0.0545
*
Gag:RNA adducts (chimera/WT)
D
gRNA
multiply spl.
7SL
n.s.
relative copies
E
RT normalized
relative copies/virus paricle
F
Other
7SL
mRNA
% reads
G
WT Total Cell
US (gRNA)
PS
MS
7SL
H
Top
Bottom
WT
Gag-SRSF5 intermediates
CA (p24)
I
Relative CA intensity
Fraction No.
J
RT activity
K
WT+NVP
Gag-SRSF5
relative copies
ERT
LRT
2-LTR
Alu-PCR


**Figure S1**

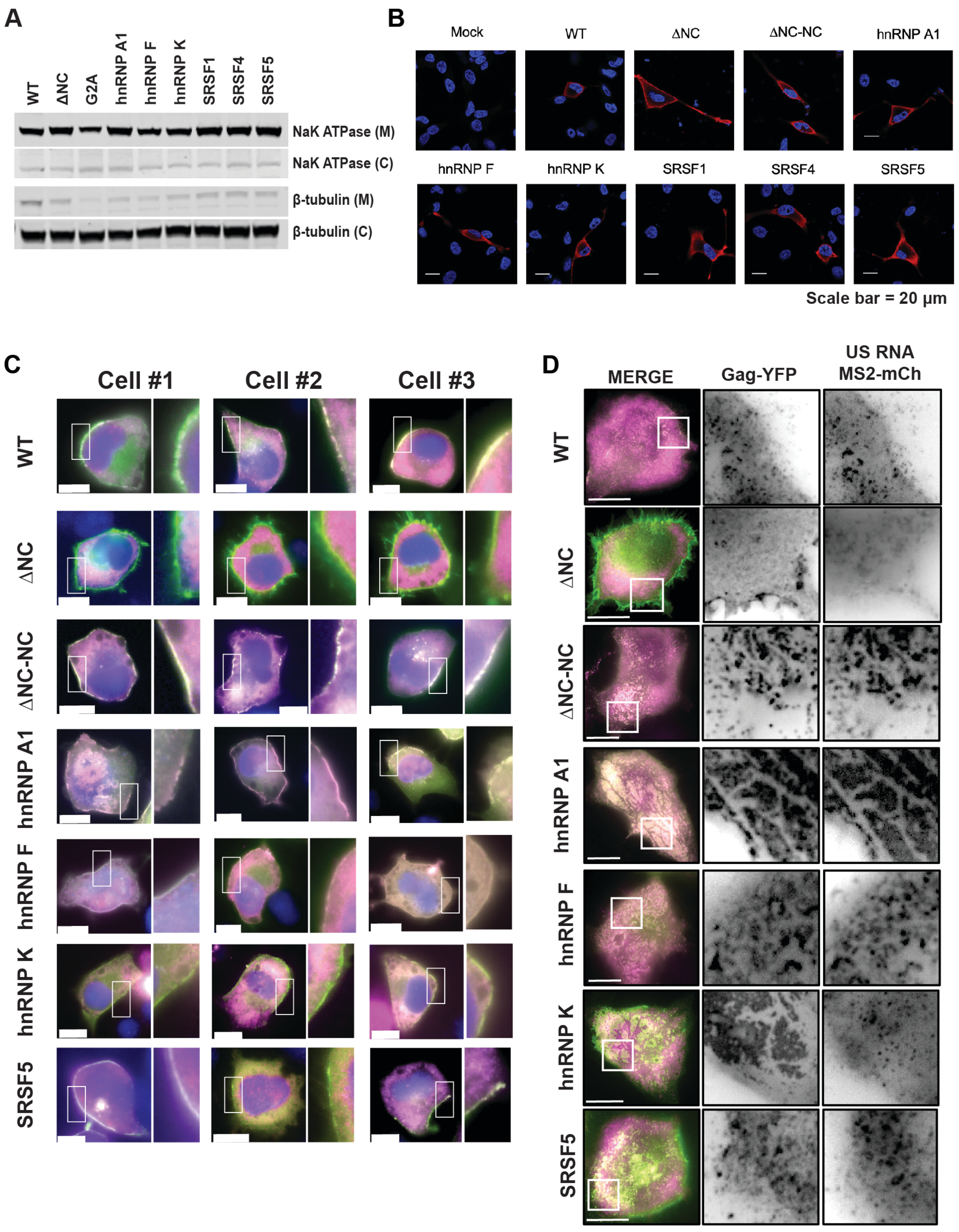

A
WT
ΔNC
G2A
hnRNP A1
hnRNP F
hnRNP K
SRSF1
SRSF4
SRSF5
NaK ATPase (M)
NaK ATPase (C)
β-tubulin (M)
β-tubulin (C)
B
Mock
WT
ΔNC
ΔNC-NC
hnRNP A1
hnRNP F
hnRNP K
SRSF1
SRSF4
SRSF5
Scale bar = 20 μm
C
Cell #1
Cell #2
Cell #3
WT
ΔNC
ΔNC-NC
hnRNP A1
hnRNP F
hnRNP K
SRSF5
D
MERGE
Gag-YFP
US RNA MS2-mCh
WT
ΔNC
ΔNC-NC
hnRNP A1
hnRNP F
hnRNP K
SRSF5


**Figure S2**

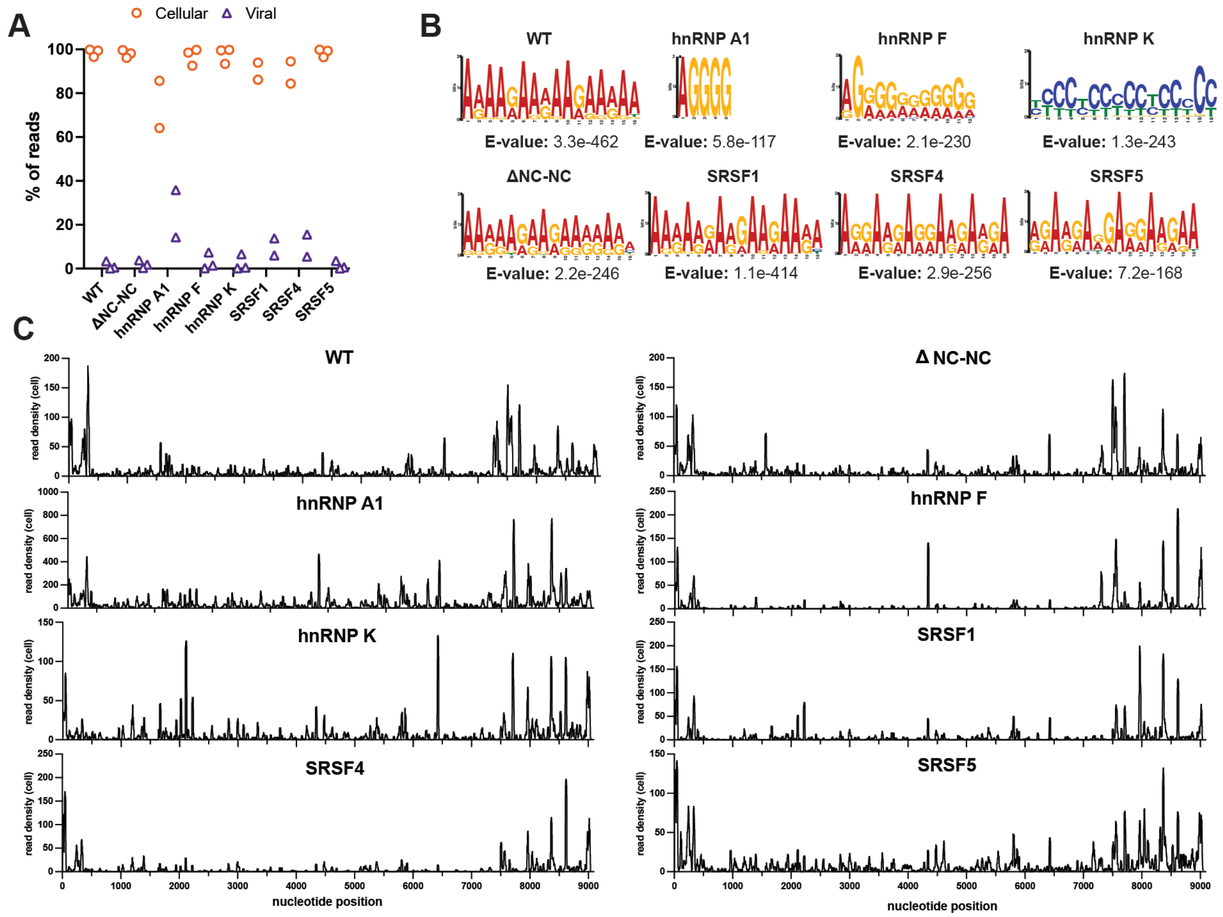


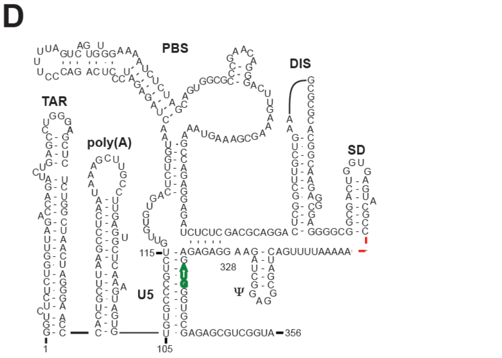


| Mutant | Deleted nucleotide positions |
|---|---|
| ΔΨ1 | Δ298-332 (ΔSL3, SD truncated) |
| ΔΨ2 | Δ301-332 (ΔSL3) |
| ΔΨ3 | Δ189-278, Δ301-332 (ΔPBS and ΔDIS) |
| ΔΨ4 | Δ105-278 (ΔU5, ΔPBS, and ΔDIS) |
| ΔΨ5 | Δ105-278, Δ298-332 (ΔU5, ΔPBS, ΔDIS, ΔSL3; SD truncated) |
| ΔΨ6 | Δ105-278, Δ301-332 (ΔU5, ΔPBS, ΔDIS, ΔSL3) |

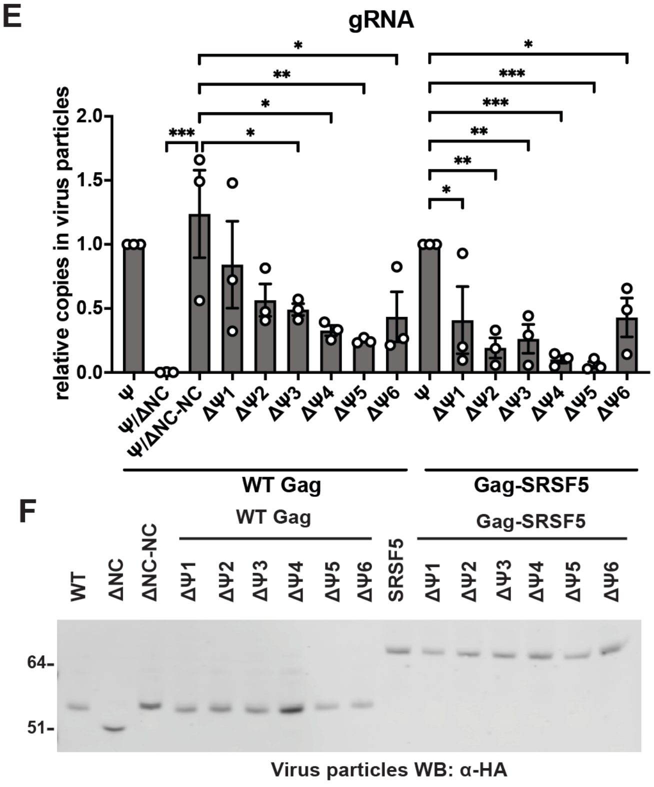


**Figure S3**

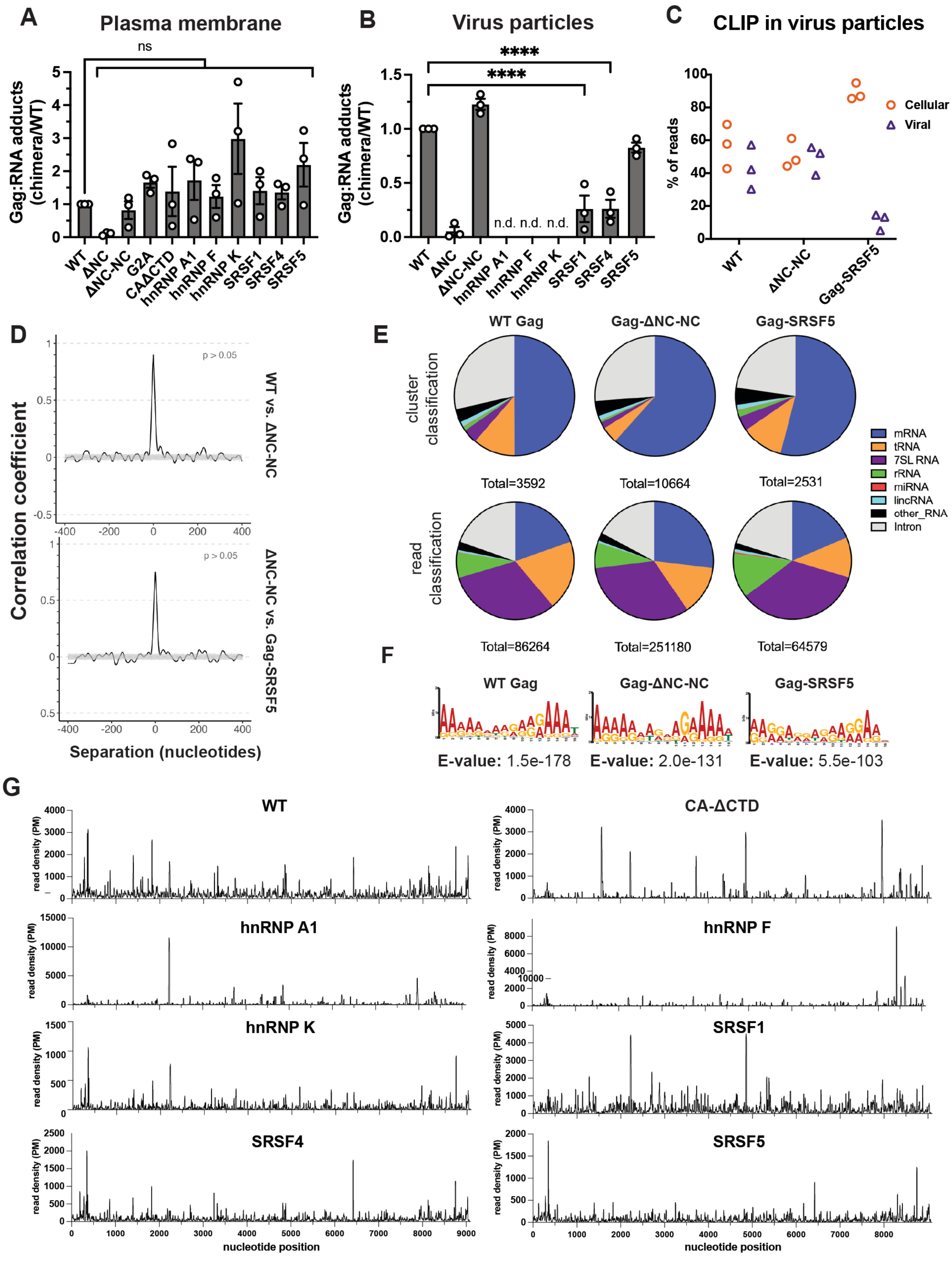
A
Plasma membrane
Gag:RNA adducts (chimera/WT)
ns
WT
ΔNC
ΔNC-NC
G2A
CAΔCTD
hnRNP A1
hnRNP F
hnRNP K
SRSF1
SRSF4
SRSF5
B
Virus particles
****
****
n.d. n.d. n.d.
C
CLIP in virus particles
% of reads
Cellular
Viral
Gag-SRSF5
D
Correlation coefficient
p > 0.05
WT vs. ΔNC-NC
ΔNC-NC vs. Gag-SRSF5
Separation (nucleotides)
E
WT Gag
Gag-ΔNC-NC
Gag-SRSF5
cluster classification
Total=3592
Total=10664
Total=2531
read classification
Total=86264
Total=251180
Total=64579
mRNA
tRNA
7SL RNA
rRNA
miRNA
lincRNA
other_RNA
Intron
F
E-value: 1.5e-178
E-value: 2.0e-131
E-value: 5.5e-103
G
WT
CA-ΔCTD
hnRNP A1
hnRNP F
hnRNP K
SRSF1
SRSF4
SRSF5
read density (PM)
nucleotide position

**Figure S4**

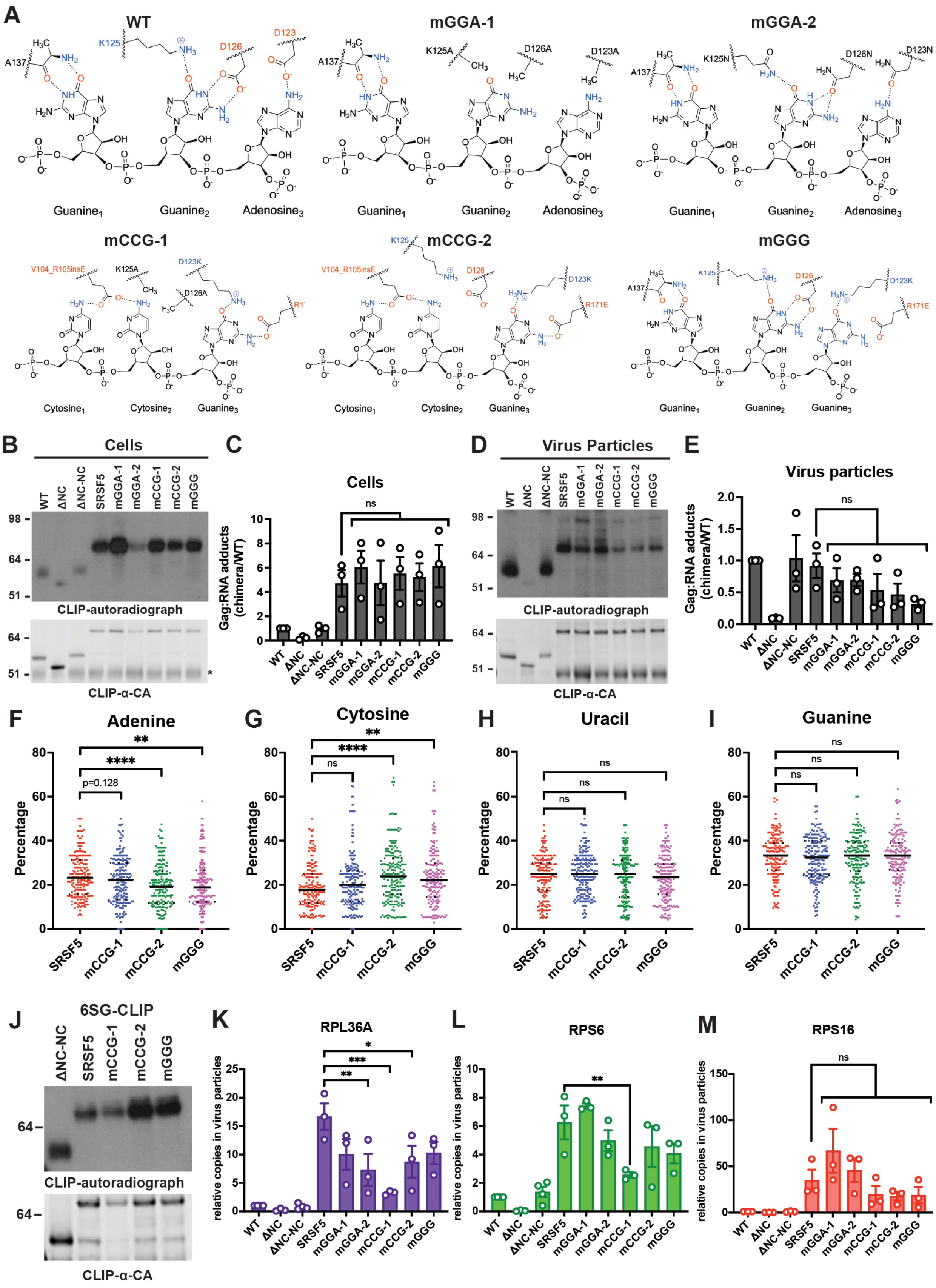

A
WT
mGGA-1
mGGA-2
Guanine1 Guanine2 Adenosine3
mCCG-1
mCCG-2
mGGG
Cytosine1 Cytosine2 Guanine3
Guanine1 Guanine2 Guanine3
B
Cells
WT ΔNC ΔNC-NC SRSF5 mGGA-1 mGGA-2 mCCG-1 mCCG-2 mGGG
CLIP-autoradiograph
CLIP-α-CA
C
Cells
Gag:RNA adducts (chimera/WT)
D
Virus Particles
E
Virus particles
F
Adenine
G
Cytosine
H
Uracil
I
Guanine
Percentage
SRSF5 mCCG-1 mCCG-2 mGGG
p=0.128
J
6SG-CLIP
ΔNC-NC SRSF5 mCCG-1 mCCG-2 mGGG
K
RPL36A
L
RPS6
M
RPS16
relative copies in virus particles


**Figure S5**

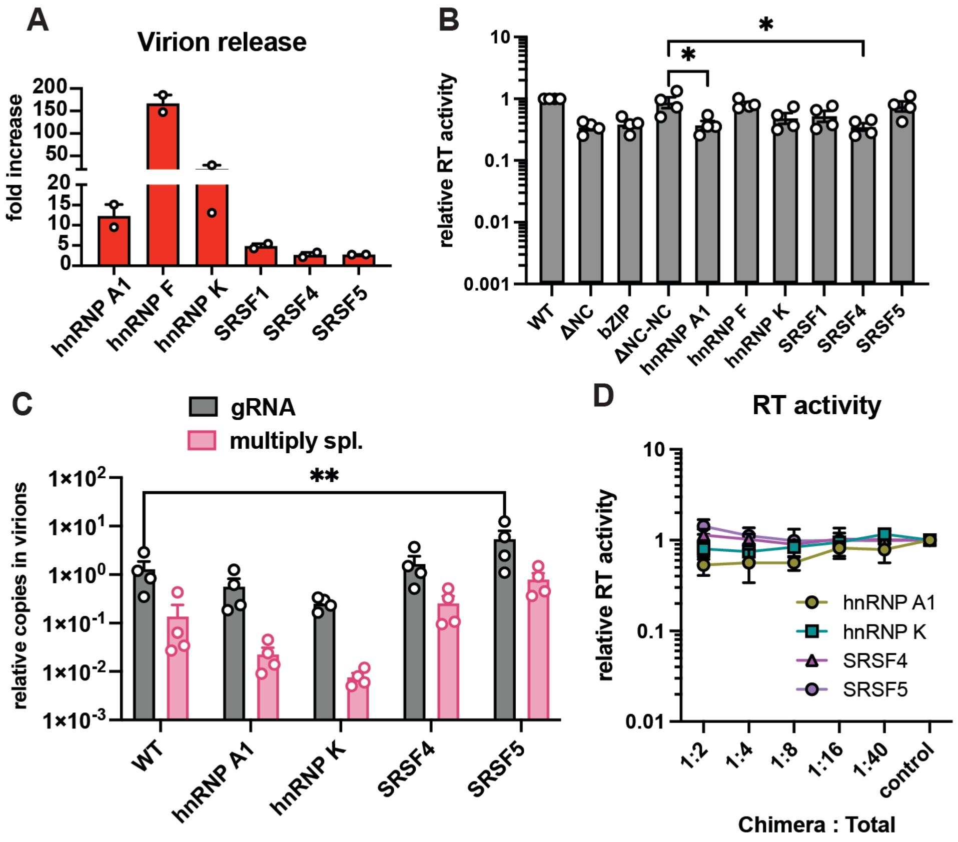
A
Virion release
fold increase
200
150
100
50
20
15
10
5
0
hnRNP A1
hnRNP F
hnRNP K
SRSF1
SRSF4
SRSF5
B
relative RT activity
10
1
0.1
0.01
0.001
*
*
WT
ΔNC
bZIP
ΔNC-NC
hnRNP A1
hnRNP F
hnRNP K
SRSF1
SRSF4
SRSF5
C
gRNA
multiply spl.
relative copies in virions
**
WT
hnRNP A1
hnRNP K
SRSF4
SRSF5
D
RT activity
relative RT activity
10
1
0.1
0.01
hnRNP A1
hnRNP K
SRSF4
SRSF5
1:2
1:4
1:8
1:16
1:40
control
Chimera : Total

**Figure S6**

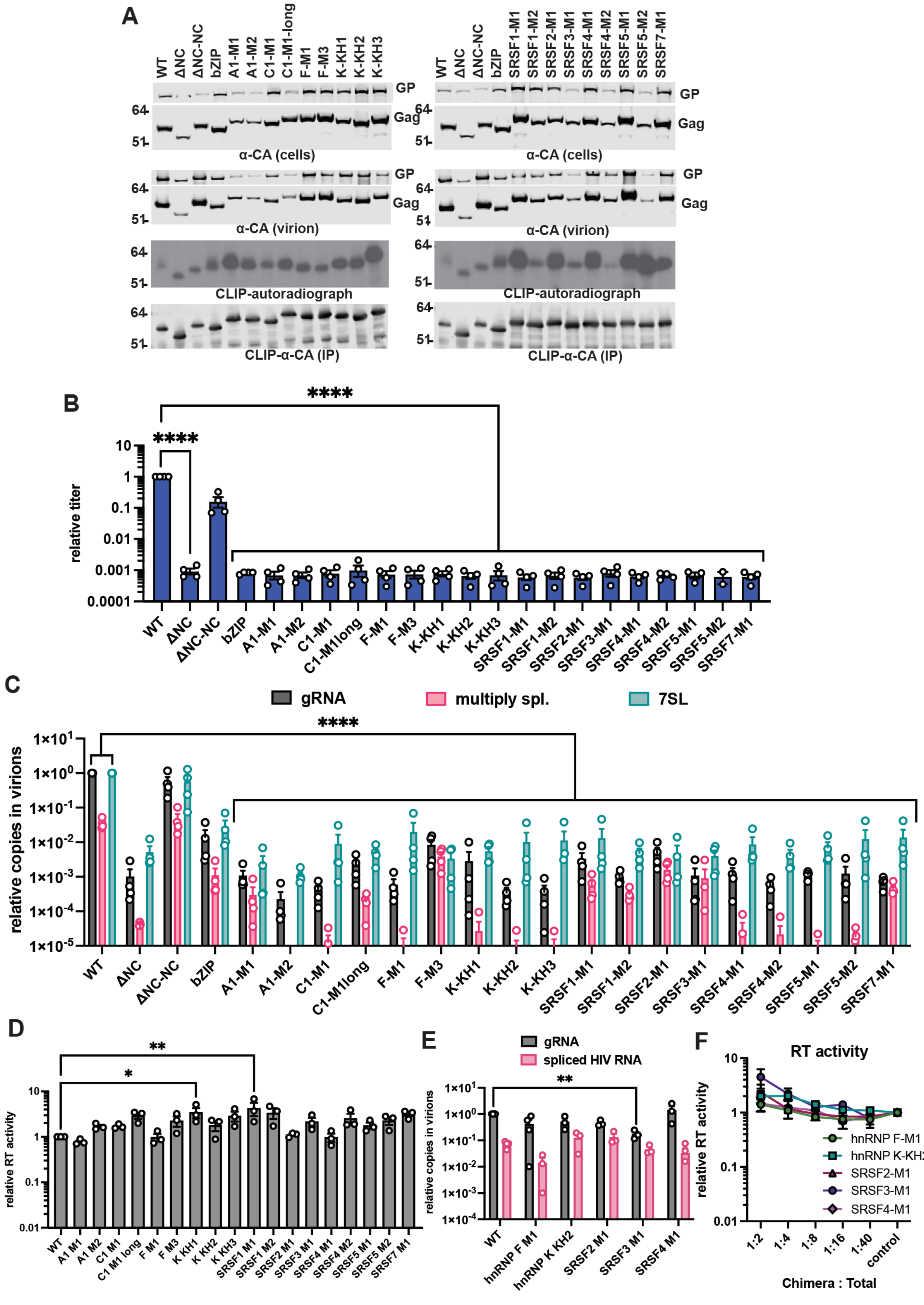

A
WT
ΔNC
ΔNC-NC
bZIP
A1-M1
A1-M2
C1-M1
C1-M1-long
F-M1
F-M3
K-KH1
K-KH2
K-KH3
SRSF1-M1
SRSF1-M2
SRSF2-M1
SRSF3-M1
SRSF4-M1
SRSF4-M2
SRSF5-M1
SRSF5-M2
SRSF7-M1
GP
Gag
α-CA (cells)
α-CA (virion)
CLIP-autoradiograph
CLIP-α-CA (IP)
B
relative titer
C
gRNA
multiply spl.
7SL
relative copies in virions
D
relative RT activity
E
spliced HIV RNA
hnRNP F M1
hnRNP K KH2
SRSF2 M1
SRSF3 M1
SRSF4 M1
F
RT activity
hnRNP F-M1
hnRNP K-KH2
SRSF2-M1
SRSF3-M1
SRSF4-M1
Chimera : Total
control


Figure S7
